# Transcriptomic profiling uncovers novel players in innate immunity in *Arabidopsis thaliana*

**DOI:** 10.1101/2021.01.02.425067

**Authors:** Mehdi Safaeizadeh, Thomas Boller, Claude Becker

**Author notes:** Author contact information: To whom correspondence should be addressed: Mehdi Safaeizadeh. Author contact information: First author: Assistant Prof. Dr. Mehdi Safaeizadeh, **E-mail:**, Contact number: Tel: +41(0)76 / 617-0006; +98(0)91 / 2037-3610; +98(0)21 / 2243-1964, Second author: Prof. em. Dr. Thomas Boller, **E-mail:**, Contact number, Third author: Prof. Dr. Claude Becker, **E-mail:**, Contact number. **Reference numbers:** The raw sequencing reads have been deposited in the EBI Annotate repository (https://www.ebi.ac.uk/fg/annotare/) under the accession number E-MTAB-9838.

## Abstract

In this research a high-throughput RNA sequencing based transcriptome analysis technique (RNA-Seq) was used to evaluate differentially expressed genes (DEGs) in the wild type Arabidopsis seedling in response to flg22, a well-known microbe-associated molecular pattern (MAMP), and *At*Pep1, a well-known peptide representing an endogenous damage-associated molecular patterns (DAMP). The results of our study revealed that 1895 (1634 up-regulated and 261 down-regulated) and 2271 (1706 up-regulated and 565 down-regulated) significant differentially expressed genes in response to flg22 and *At*Pep1 treatment, respectively. Among significant DEGs, we observed that a number of hitherto overlooked genes have been found to be induced upon treatment with either flg22 or with *At*Pep1, indicating their possible involvement in innate immunity. Here, we characterized two of them, namely PP2-B13 and ACLP1. *pp2-b13* and *aclp1* mutants showed an increased susceptibility to infection by the virulent pathogen *Pseudomomas syringae pv tomato mutant hrcC-*, as evidenced by an increased growth of the pathogen in planta. Further we present evidence that the *aclp1* mutant was deficient in ethylene production upon flg22 treatment, while the *pp2-b13* mutant, was deficient in ROS production. The results from this research provide new information to a better understanding of the immune system in Arabidopsis.

## Introduction

As sessile organisms, plants are constantly under attack by a broad range of different microbes^1–7^. In a co-evolutionary arms race between plants and pathogens, plants initially sense the presence of microbes by perceiving microbe-associated molecular patterns (MAMPs) via membrane-resident pattern recognition receptors (PRRs) that are located on the cell surface; such MAMP perception generally leads to pattern-triggered immunity (PTI) ^1,4,8,10^.

The model plant *Arabidosis thaliana* can detect a variety of MAMPs, including fungal chitin and bacterial elicitors such as flagellin and elongation factor-Tu (EF-Tu), or their respective peptide surrogates flg22 and elf18^8–10^. Flagellin and EF-Tu are perceived by FLS2 and EFR receptors, respectively. Besides MAMPs, molecular patterns derived from the plant upon pathogen attack can also trigger an immunity response. Examples of such damage-associated molecular patterns (DAMPs) are members of the family of AtPeps, recently discovered endogenous and highly conserved peptides in *A. thaliana*. The different AtPeps (AtPep1-8) originate from the conserved C-terminal portion of their respective precursors AtPROPEP1–8^11–14^. The plant cell surface PRRs *At*PEPR1 and *At*PEPR2 have been identified as the AtPeps receptors^12,16,17^.

DAMP/MAMP perception triggers a vast array of defense responses^1,12^. These include the production of reactive oxygen species (ROS) in an oxidative burst^18,19^, the multi-level specific reprogramming of expression profiles at transcriptional and also post-transcriptional levels^20–23^, and downstream defense responses, including callose deposition^24^, MAP kinase activation, synthesis of the defense hormone salicylic acid (SA), and seedling growth inhibition^25^. MAMP treatment prior to the actual pathogen attack results in enhanced resistance to adapted pathogens, and it has been observed that mutants impaired in MAMP recognition display enhanced susceptibility, not only to adapted but also to non-adapted pathogens^10,19,23,26^. This indicates a contribution of pattern-triggered immunity (PTI) to both basal and non-host resistance, highlighting the importance of PTI in plant innate immunity^27–30^.

The proteobacterium *Pseudomonas syringae* is a bacterial leaf pathogen that causes destructive chlorosis and necrotic spots in different plant species, including monocots and dicots. *P. syringae* pathovars and races differ in host range among crop species and cultivars, respectively^6,31^. Many strains of *P. syringae* are pathogenic in the model plant *A. thaliana*, which makes *P. syringae* an ideal model to investigate plant–pathogen interactions^31–33^. The ability of *P. syringae* to grow in plants and to multiply endophytically depends on the type-three secretion system (T3SS). T3SS enables the secretion into the cytoplasm of the plant cell of effector proteins, which suppress or, in some cases, change plant defense responses^34^. *P. syringae* encodes 57 families of different effectors injected into the plant cell by the T3SS^32^. Effectors inside plant cells are recognized by R proteins, which constitute the second level of defense known as effector-triggered immunity (ETI)^1,20,35^.

PTI response is controlled by a complex, interconnected signaling network, including many transcription factors (TFs); interference with this network can paralyze the adequate response upon pathogen infection^36,37^. A large fraction of genes in the plant genome respond transcriptionally to pathogen attack^21,38^. In addition to specific reprogramming of transcription, post-transcriptional regulation also plays a role in the plant immune response^39^. The advent of advanced sequencing and proteomics technologies has led to the identification of many novel players in defense signaling pathways and their characterization as important components of innate immunity in *Arabidopsis*. However, for a fundamental understanding of the plant’s defense system and its response to pathogens, it is necessary to fill the remaining gaps by further identifying genes and proteins involved in plant immunity^1^.

The highly conserved 22-amino-acid fragment (flg22) of bacterial flagellin that is recognized by the FLS2 PRR can activate an array of immune responses in Arabidopsis^1–4^. In addition, resistance to *Pst* DC3000 is induced by pre-treatment with flg22^1–4,9,23^. Previous studies investigating flg22-induced transcriptional changes showed that among highly induced genes, there were several with functions in the Arabidopsis immune pathway that had previously not been associated with immunity^23,40,41^. These studies profiled only a part of the Arabidopsis gene space, and one can therefore speculate that a whole-genome transcriptome profiling of elicitor-treated Arabidopsis plants would unveil additional new players in the immune signaling system. Furthermore, given that both the MAMP flg22 and the DAMPs AtPeps trigger immunity, analyzing their respective effects side by side in one coherently designed experiment could increase the power to detect shared features and specific responses of the respective immune response pathways.

Here, we performed whole-genome transcriptome profiling by RNA sequencing (RNA-seq)^42–45^ of Arabidopsis seedlings treated with either flg22 or AtPep1 treatments. Filtering for genes induced in both treatments and those missing in previously published assays, we selected 85 candidate genes to be investigated for their role in plant immune response and systematically tested T-DNA insertion mutants of these genes for susceptibility towards *Pst*. For two loci, *PHLOEM PROTEIN 2-B13* (*PP2-B13*) and *ACTIN CROSS-LINKING PROTEIN 1* (*ACLP1*), we identified mutant lines with altered pathogen response phenotypes and characterized these genes as novel players in early PTI responses.

## Results

### Whole-genome transcriptional profiling identifies two novel factors of PTI

To dissect transcriptional responses in response to flg22 and AtPep1, we extracted total RNA from mock- and elicitor-treated one-week-old Arabidopsis plants and performed RNA-seq transcriptome analysis on three biological replicates per treatment (Supplementary File S1 and S2). Samples were collected 30 min after elicitor treatment. We used the R package DESeq2^46^ for differential gene expression analysis; all differentially expressed genes (DEGs) can be found in Supplementary Files S3-S6. In response to flg22, we detected a total of 1,895 DEGs compared to the control treatment (Fig. 1A), of which 1,634 genes were up- and 261 were down-regulated in the flg22-treated seedlings (Supplementary Files S3 and S5). Treatment with AtPep1 resulted in 2,271 DEGs, with 1,706 up-regulated and 565 down-regulated (Fig. 1A). When comparing the two treatments with each other, we detected only 511 DEGs, with similar fractions of up- and down-regulated genes (265 and 246, respectively, in flg22 vs. AtPep1) (Fig. 1A). Taken together, these results indicated that AtPep1 treatment causes slightly more genes to be differentially regulated than flg22, and that the transcriptional profiles are more similar between flg22- and AtPep1-treated samples than between either of the treatments and the control. While a remarkable 70% of flg22-up-regulated genes were also induced by AtPep1, 256 genes were exclusively up-regulated in response to flg22, while 328 were exclusively up-regulated in response to AtPep1 (Fig. 1; panel B). Of genes down-regulated upon flg22 treatment, only 23% were also down-regulated in response to AtPep1; 107 genes were exclusively down-regulated by flg22 treatment, vs. 411 genes by AtPep1 (Fig. 1; panel C). Detailed information on DEGs from different comparisons are presented in Supplementary Files S1 to S16.

**Figure 1.**
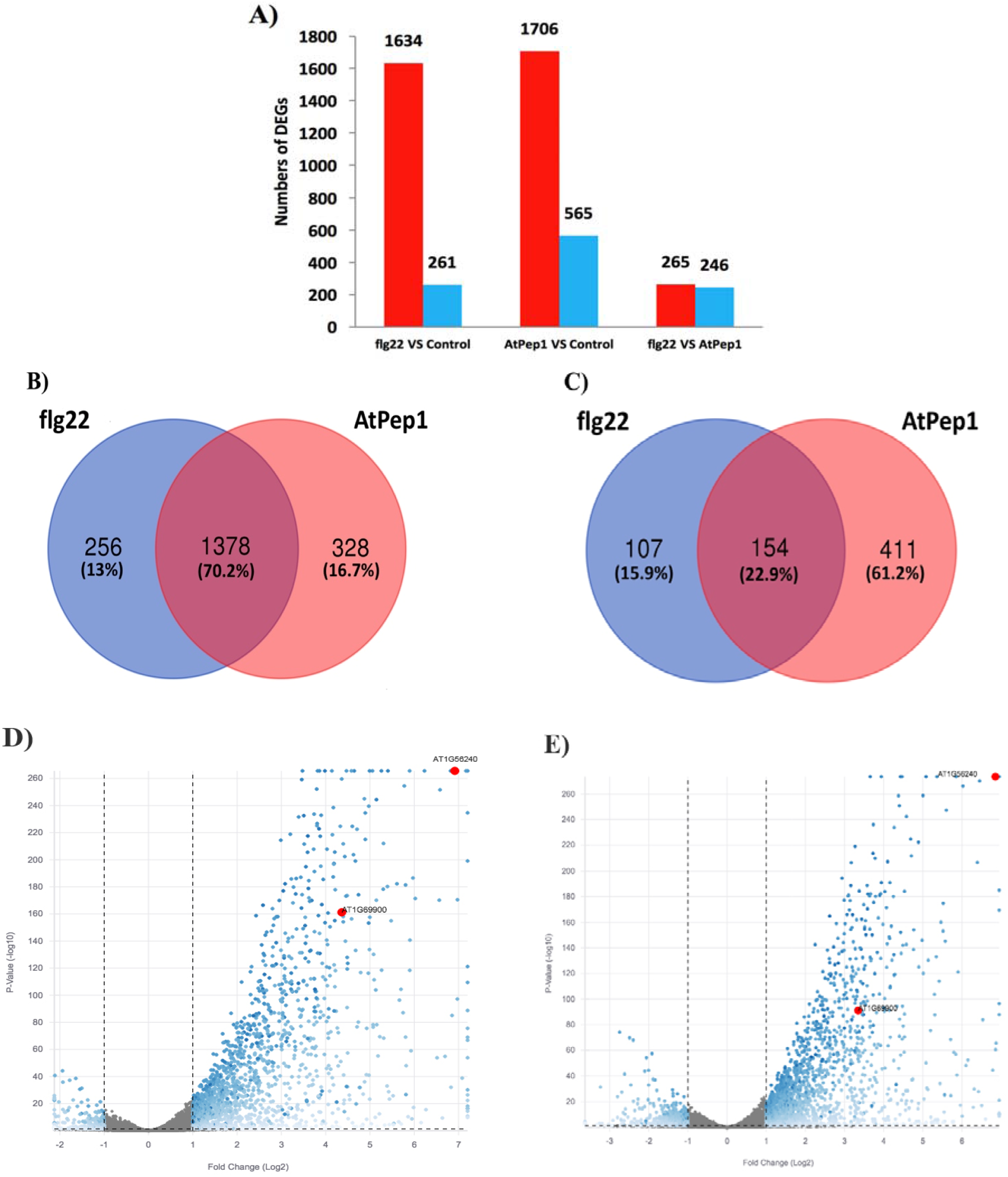
Distribution of differentially expressed genes (DEGs). (A) DEGs in Arabidopsis thaliana in response to in in response to flg22 and AtPep1 treatments compared to the control investigated in this study. (**B**) Venn diagram of up-regulated DEGs between flg22 treatment and AtPep1 treatments. (**C**) Venn diagram of down-regulated DEGs between flg22 treatment and AtPep1 treatments. The overlapping regions display the common transcripts. (**D**) Volcano plot of DEGs in response to flg22 treatment; (**E**) Volcano plot of DEGs in response to AtPep1 treatment. In (**D**-**E**), blue dots correspond to significantly up- and down-regulated DEGs, grey dots represent non-DEGs. *At1G56240* (*PP2-B13*) and *At1G69900* (*ACLP1*) are highlighted in red.

In response to flg22, the expression levels of *PP2-B13* and of *ACPL1* were 126-fold and 20-fold induced, respectively (Fig. 1D). Similarly, AtPep1 treatment leads to 120-fold up-regulation of *PP2-B13* and a 10-fold up-regulation of *ACLP1* (Fig. 1E).

Former studies showed that treatment of Arabidopsis seedlings with flg22 triggers robust PTI-like responses at the transcriptional level, activating ca. 1,000 genes that may have functions in PTI responses^23,40,41^. However, because these experiments were done using the ATH1 microarray, which does not cover all Arabidopsis protein-coding genes, we speculated that there might be additional, so far unknown PTI-related genes affected by flg22 and other elicitors. Denoux et al., (2008)^41^ performed a comprehensive microarray (Affymetrix ATH1) transcript analysis in response to flg22 treatment. However, our RNA-seq analysis revealed 3,297 genes induced (without fold-change cutoff) in response to flg22 treatment that had not been present on the ATH1 chip (Supplementary File S7). Comparing the upregulated DEGs results in RNA-seq experiment analysis with fold change cutoff (adjusted *p*-value < 0.05 and a minimum two-fold change) among the genes which are also present in ATH1 affymetrix genechip showed that 1366 upregulated DEGs are present in both RNA-seq experiment and ATH1 affymetirx genechip (Supplementary File S8). While our analysis showed that 268 genes with fold change cutoff are exclusively upregulated in RNA-seq analysis which were not present in ATH1 affymetrix genechip and their expression only investigated in RNA-seq analysis (Supplementary File S9). To identify yet unknown PTI factors, we first discarded all genes from our list of DEGs that had been present on the ATH1 microarray chip and hence would have been detected in the above-mentioned studies.

We then ranked the remaining DEGs by fold change induction (high induction of transcription in response to both flg22 and AtPep1 treatments) and selected the 85 most strongly up-regulated genes as candidates (Supplementary File S10). Finally, we decided to focus on a small set of genes that showed highest induction after flg22 treatment (Table 1). We checked for availability of T-DNA insertion mutants for these genes and retrieved mutant lines for AT1G56240, AT1G65385, AT4G23215, AT1G59865, AT1G24145, AT2G35658, AT1G69900, AT2G27389, and AT1G30755. We confirmed homozygous T-DNA insertions via PCR.

**Table 1.**
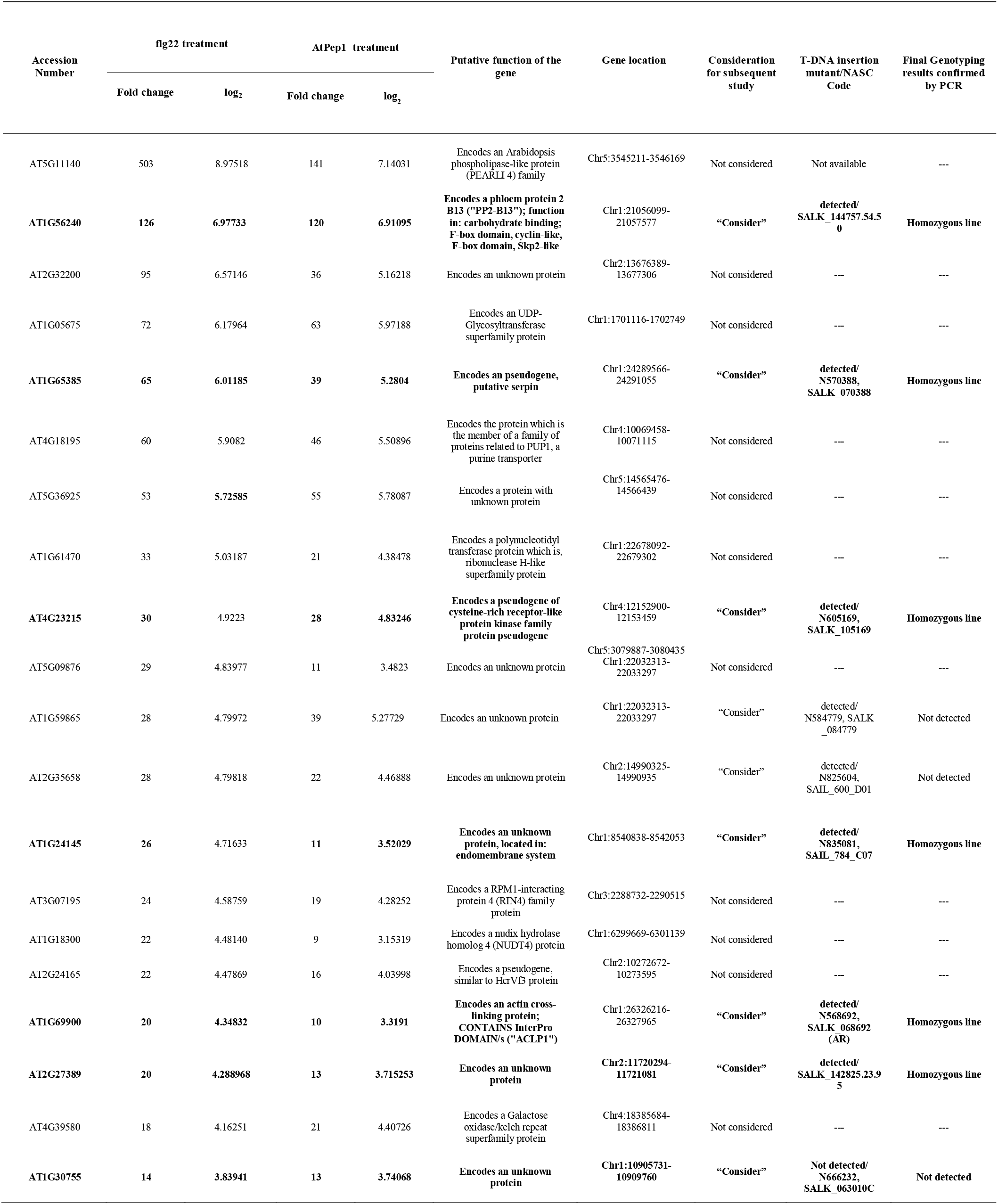
Detailed information on the top up-regulated genes based on RNA-seq analysis (fold change 30 minutes after flg22 and AtPep1 treatments compared to the control). Genes of interest are highlighted in bold.

To test whether any of the candidate genes might play a role in immunity, we tested all homozygous T-DNA mutant lines for bacterial growth of the mutant pathogenic strain *P. syringae* pv. *tomato hrcC-* (*Pst hrcC-*), which is defective in T3SS. Comparing the bacterial growth titer in the mutant plants to that of wild-type Col-0 revealed that two of the lines, namely SALK_144757.54.50 and SALK_68692.47.55, showed significantly better bacterial growth (*P*-value = 0.0261 and 0.0089, respectively; Student’s *T*-test) (Fig. 4), and that the underlying loci might play a role in defense signaling.

### Expression of the *PP2-B13* and *ACLP1* genes is induced following flg22 treatment

According to the Arabidopsis Information Resource^47^ and the SIGnAL database (http://signal.salk.edu/), the predicted T-DNA insertion site in SALK_144757.54.50 is located in the second of three exons of *PP2-B13*^48^ (Fig. 3A); the T-DNA insertion in SALK_68692.47.55 is located in the first of two exons of *ACLP1*. We confirmed that the T-DNA insertion lines were null alleles for *pp2-b13* and *aclp1*, respectively, via reverse-transcription polymerase chain reaction (RT-PCR) (Fig. 3B). *PP2-B13* and *ACLP1* transcripts were not detectable in the respective T-DNA insertion lines; we therefore refer to these lines as *pp2-b13* and *aclp1*, respectively. Visual inspection of plant growth did not reveal any obvious phenotypic differences between any of the two insertion lines and wild-type Col-0 with regard to size and shape at the rosette stage. Two days post infection with *Pst hrcC-*, neither *pp2-b13* nor *aclp1* showed symptoms different to this of with wild-type Arabidopsis (Supplementary Fig. S4).

As can be seen in the volcano plot in Figure 1D, gene expression levels of *PP2-B13* and *ACLP1* were strongly induced by flg22. To further monitor the gene expression of *PP2-B13* and *ACLP1* upon elicitor perception and to validate the RNA-seq results, we analyzed expression levels by quantitative real-time PCR (qRT-PCR) in leaves of four-week-old Arabidopsis plants at different time points. We confirmed that also at this later developmental stage, expression of *PP2-B13* and *ACLP1* was strongly induced (100-fold for *PP2-B13* and 12-fold for *ACLP1*) within 30 minutes after flg22 treatment (Fig. 2). Two and six hours after elicitor treatment, expression levels of *PP2-B13* had returned to pre-treatment levels, while those of *ACLP1* remained only slightly elevated (Fig. 2). This expression pattern suggests that both genes might be involved in early defense response.

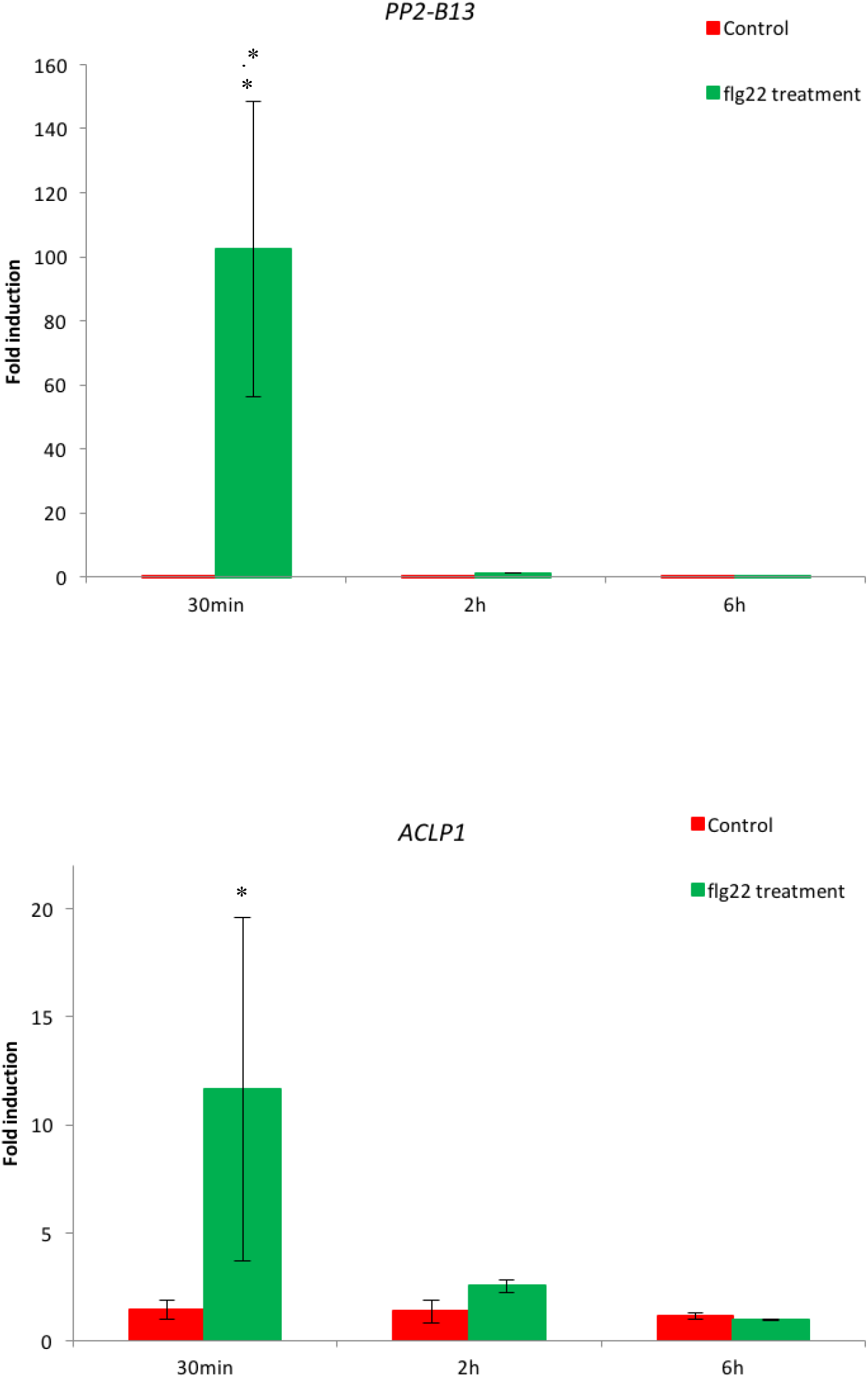

**Figure 3.**
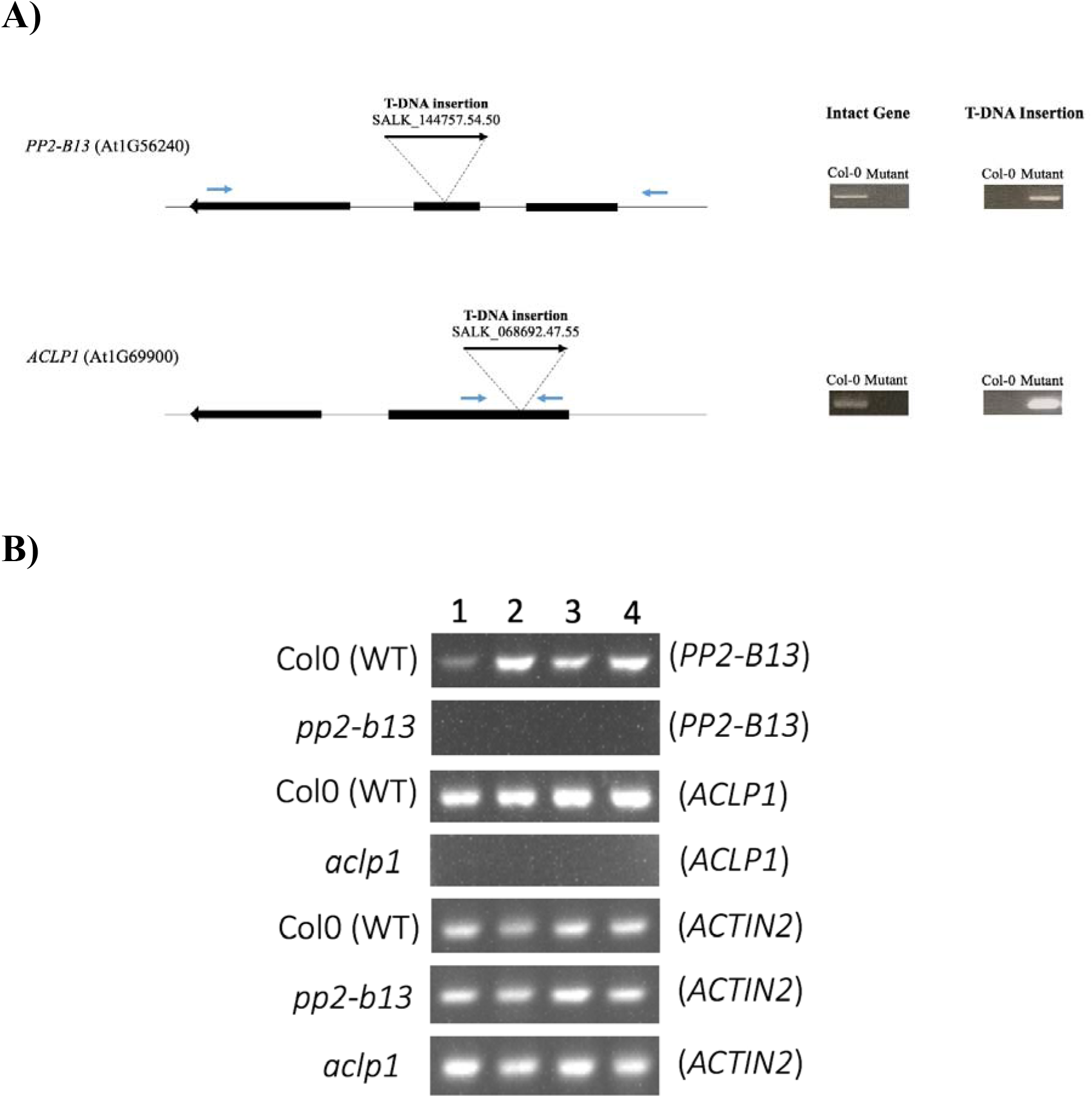
A) Schematic representation of homozygous T-DNA mutant lines *PP2-B13*, and *ACLP1*. Boxes indicate exons; thin lines indicate introns; bold arrows indicate T-DNA insertions; arrows indicate the direction of the gene. On the right side, the PCR results of the homozygous lines are shown, amplifying either the intact gene or the T-DNA. Small blue arrows indicate the primers position. **B**) RT-PCR results showing transcripts in Col-0 (WT), *pp2-b13* and *aclp1* mutant lines. The lower panel shows amplification of *ACTIN2* transcript as a control. Numbers 1 to 4 indicate individual plants for each genotype.

**Figure 4.**
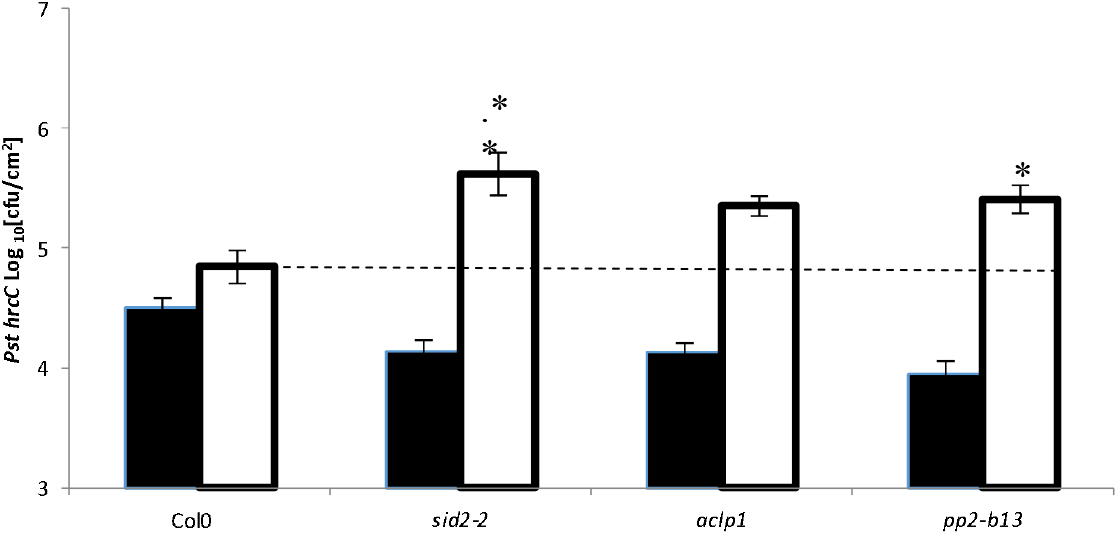
Bacterial susceptibility assay. Leaves of four-to-six-week-old Arabidopsis plants (Col-0, *sid2-2*, *pp2-b13*, and *aclp1*) were pressure infiltrated with *Pseudomonas syringae* pv. *tomato* mutant *hrc*C-(OD_600_=0.0002, in infiltration buffer). *sid2-2* mutant plants, which are deficient in salicylic acid production, were used as a positive control. Black bars indicate bacterial colony from leaf discs of infected leaves just after infiltration (0 day); white bars represent colony-forming units (cfu/cm2) 48 h post inoculation. Bars show the mean ± s.e. (n=6). Similar results were observed in four independent experiments. Asterisks indicate a significant difference (*p ≤0.05, **p≤0.01) from the wild type plants as determined by Student’s *t-test*.

### Increased susceptibility to *Pseudomonas syringae* pv. *tomato* mutant *hrc*C- in *pp2-b13* and *aclp1* mutant lines

Two days post inoculation of leaves with *Pst hrcC-*, the bacterial titer for wild type Arabidopsis reached 109,000 cfu/cm^2^, while for *pp2-b13* mutant lines it increased significantly to 325,000 cfu/cm^2^ (*p* = *0*.0261), albeit not as drastically as that of *sid2-2* mutants. The protein encoded by *PP2-B13* is a phloem protein containing the F-box domain Skp2. It also has a described function in carbohydrate binding^48^.

The protein encoded by *ACLP1* is of unknown function with the highest similarity to actin cross-linking proteins and includes a fascin domain. As it can be seen in Fig. 4, 48 hours post inoculation of leaves with *Pst hrcC-*, the bacterial titer for wild type Arabidopsis, reached 109000 cfu/cm^2^ while for the *aclp1* mutant line, it increased significantly to 257000 cfu/cm^2^ (p = 0.0089).

In conclusion, these results suggest that PP2-B13, and ACLP1 play a role in defense signaling and that both genes are required for wild-type levels of resistance against *Pst hrcC-*.

### Differential ethylene production in *pp2-b13* and *aclp1* plants, as compared to the wild type Arabidopsis

To analyze the early defense responses upon elicitor treatment, we assessed ethylene (ET) production in response to flg22 treatment in the mutant lines *pp2-b13*, and *aclp1.* We observed that mutant line *aclp1* displayed a significantly reduced ET production in comparison to wild-type Arabidopsis upon treatment with 1 μM flg22 (*p-*value = 0.0295; Fig. 5). This suggests that ACLP1 is involved in the enhancement of ET production in response to flg22 perception.

**Figure 5.**
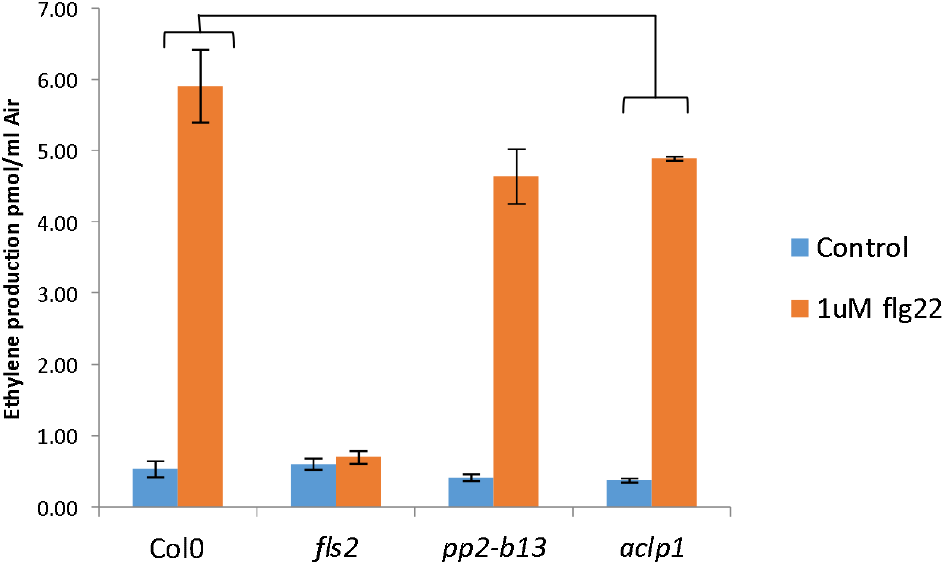
Early PTI responses upon elicitor treatment. Ethylene accumulation after elicitor treatment. Leaf discs of four- to five-week-old plants of wild-type and mutant lines (*pp2-b13*, and *aclp1*) were treated with 1 μM of the flg22 elicitor peptide or without any peptide (control). In all cases, ethylene production was measured three and half hours after closing the tubes. Ethylene accumulation in *pp2-b13* and *aclp1* mutant lines was compared to the wild type Arabidopsis. *fls2* mutant line was used as a negative control. Columns represent mean ethylene concentration of six biological replicates. Error bars indicate standard deviation with n=6. Similar results were obtained in at least six independent biological replicates. T-test was performed comparing the responses of the control treatment to the elicitor treatments; *P* values are indicated \**p*≤0.05.

### Differential Reactive Oxygen Species Generation in *pp2-b13* and *aclp1* plants, as compared to the wild type Arabidopsis

One of the early responses triggered by MAMPs and DAMPs is the production of apoplastic ROS by the Arabidopsis NADPH-oxidases RbohD and RbohF protein^19^. We observed that in the treated leaf discs upon flg22 perception, *pp2-b13* displayed a lower ROS production compared to wild-type (Fig. 6A), indicating that PP2-B13 might play a role in early PTI by enhancing the oxidative burst in response to the flg22 perception. In contrast, *aclp1*, although exhibiting deficiency in ET production upon flg22 perception, showed robust enhancement of ROS production at levels similar to that of wild-type (Fig. 6A).

**Figure 6.**
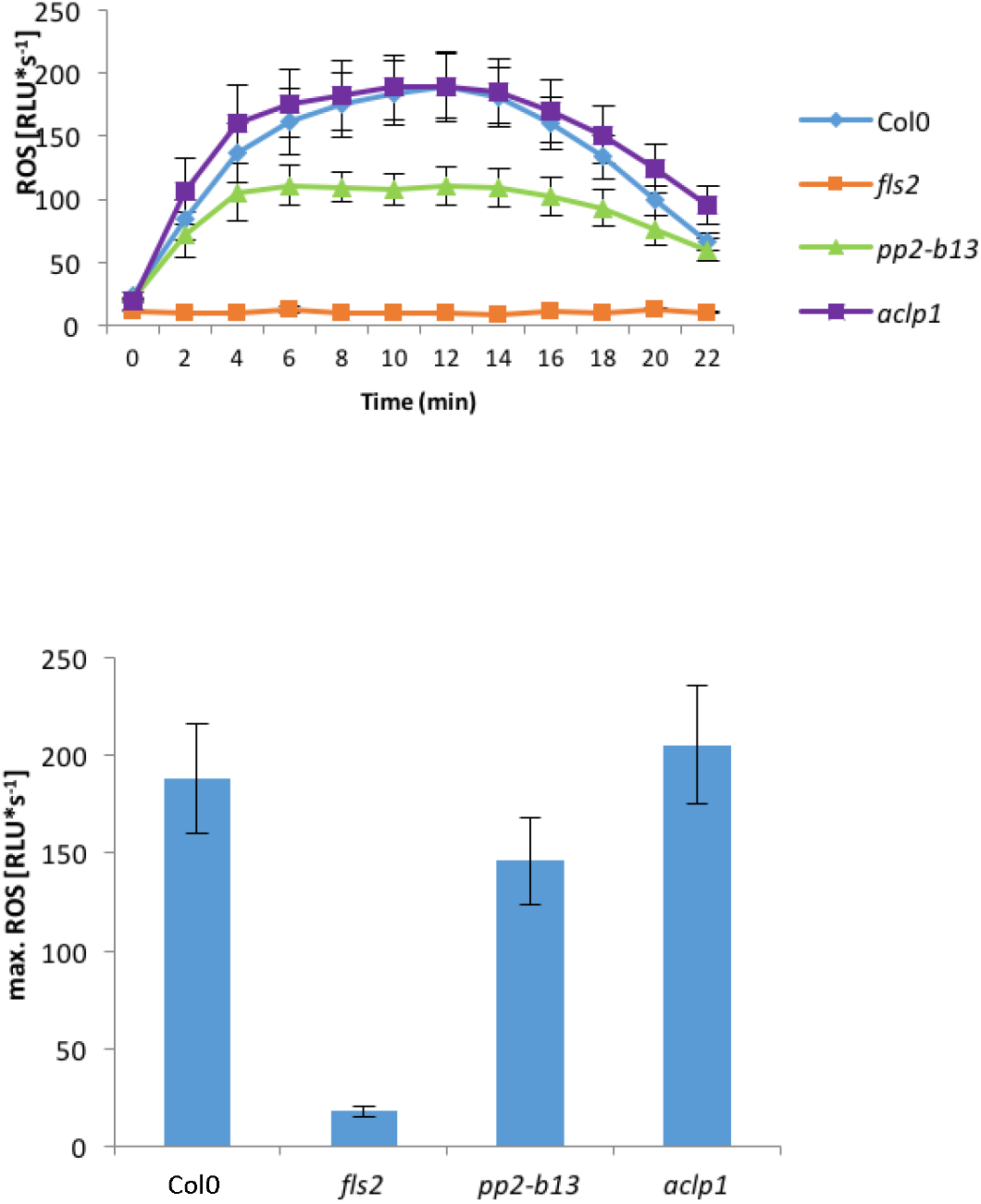
ROS production after treatment with flg22. Leaf discs were treated with indicated 1 μM flg22 orwithout any peptide (control). **(A)** indicates ROS production in *pp2-b13* and *aclp1* mutant lines compared to wild-type Arabidopsis; (**B**) represents maximum ROS production in *pp2-b13* and *aclp1* mutant lines compared to wild-type Arabidopsis. *fls2* mutant line was used as a negative control. Graphs display average of 12 replicates. Error bars indicate standard error (SE) of the mean. The experiment was repeated four times with similar results. RLU= relative light units.

When comparing maximum ROS production the two mutant lines and wild-type, we did not observe statistically significant differences, in contrast to the fls2 mutant which displayed no ROS production at all in response to flg22 (Fig. 6B).

### FLS2 receptor abundance in *pp2-b13* and *aclp1* mutants were similar to the wild type Arabidopsis

The *PP2-B13* and *ACLP1* genes were strongly induced upon elicitor treatment, as seen in the RNA-seq and qPCR data. Additionally, both mutant lines were deficient in early PTI responses (ET and ROS measurement). Hence it is conceivable that the products of the *PP2-B13* and *ACLP1* genes affect the abundance of FLS2 receptor. However, FLS2 analysis via immunoblots showed that both mutant lines had similar levels of FLS2 as the wild-type (Fig. 7), indicating that these genes do not play a role in regulating the abundance of the FLS2 receptor.

**Figure 7.**
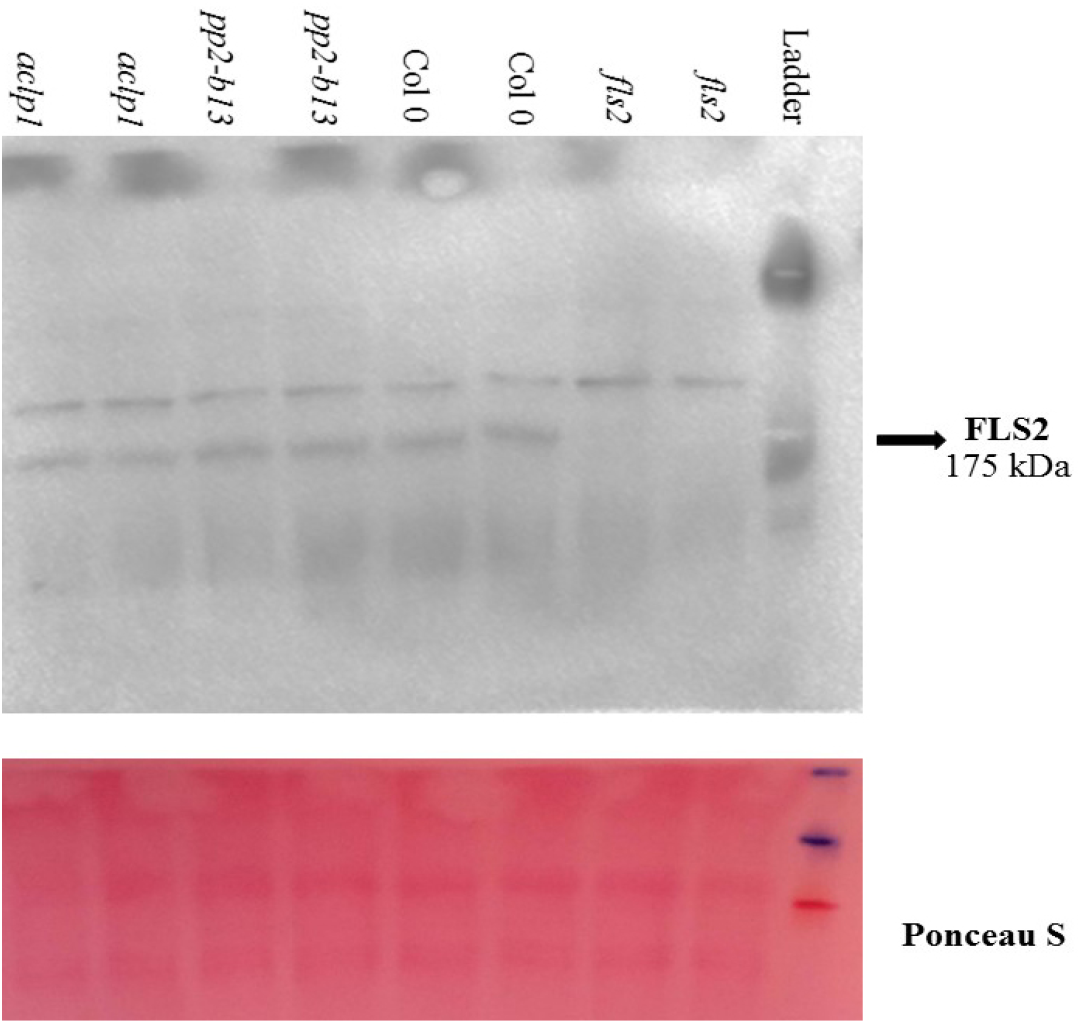
FLS2 protein levels. FLS2 protein levels of the mutant lines *pp2-b13* and *aclp1* as detected by immunoblot using a FLS2-specific antibody. *fls2* mutant plant is used as negative control. Ponceau S staining was used as loading control.

## Discussion

Plants are under constant exposure to microbial signals from potential pathogens, potential commensals, and mutualists. The plant cell immune sensors are able sense these signals and expand the defense against pathogens^1,20,49–50^. Host-pathogen interactions encompass a complex set of events that are dependent on the nature of the interacting partners, developmental stage, and environmental conditions^1,50–51^. These interactions are regulated through diverse signaling pathways that ultimately result in altered gene expression^1,23,42^. Membrane-resident pattern recognition receptors (PRRs) that are located on the cell surface can sense and perceive microbe-derived signature components known as microbe-associated molecular patterns (MAMPs) and also damage-associated molecular patterns (DAMPs), leading to pattern-triggered immunity^1–4^.

In the current study, global gene expression profiling of wild type Arabidopsis seedlings resulted in the identification of a large number of genes induced by flg22 and *At*Pep1 that had not been detected by the ATH-1 array technology. Among them, we focused on two, namely *PP2-B13*, and *ACLP1*. We observed noticeable up-regulation in wild type Arabidopsis for both of these genes upon flg22 treatment (Fig. 2). Reverse-genetic studies of *PP2-B13* and *ACLP1* genes showed that these genes are required to control infection by the bacterial pathogens *P. syringae* pv. *tomato* mutant *hrc*C-(*Pst hrcC-*; Fig. 4). Our results highlight the general usefulness of transcriptomic approaches to identify new players in early defense responses in innate immunity and reveal two new players, PP2-B13 and ACLP1, in this pathway. It should be noted that extending the time points of the elicitor treatment in future studies might help uncover additional players in innate immunity.

PP2-B13^48^ is an F-box protein with homology to PP2-B14^52^. The F-Box domain of PP2-B13 is close to the N-terminus of the protein. PP2-B13 shows the highest similarity in amino acid sequence with AT1G56250, which formerly was reported as an F-box protein^52^. Zhang *et al*., (2011)^53^ showed that PP2-B13 and PP2-B14 were highly abundant in phloem upon aphid infection. These genes are located in a cluster of defense-related genes, which supports that hypothesis that they play a role in the defense signaling network.

Sequence alignment of PP2-B13 with homologues from other plant species revealed conserved features (Fig. 8). A phylogenetic analysis supported high conservation of PP2-B13-like proteins across different plant species, suggesting similar function (Supplementary Fig. S7). *In silico* structural analysis using Raptor X^54^ predicted two domains (Supplementary Fig. S5): an N-terminal F-box domain (residues 4-46; Fig. 8) and a C-terminal PP2 domain (residues 93-280; Fig. 8). PP2-domain proteins are one of the most abundant and enigmatic proteins in the phloem sap of higher plants^55–56^. It was reported that lectin domain proteins are important in plant defense responses, and so far 10 membrane-bound lectin type PRRs, which are involved in plant defense signaling and symbiosis, have been identified^56^. Recently, Eggermont et al. (2017)^57^ showed that lectins are linked to other protein domains which are identified to have a role in stress signaling and defense.

**Figure 8.**
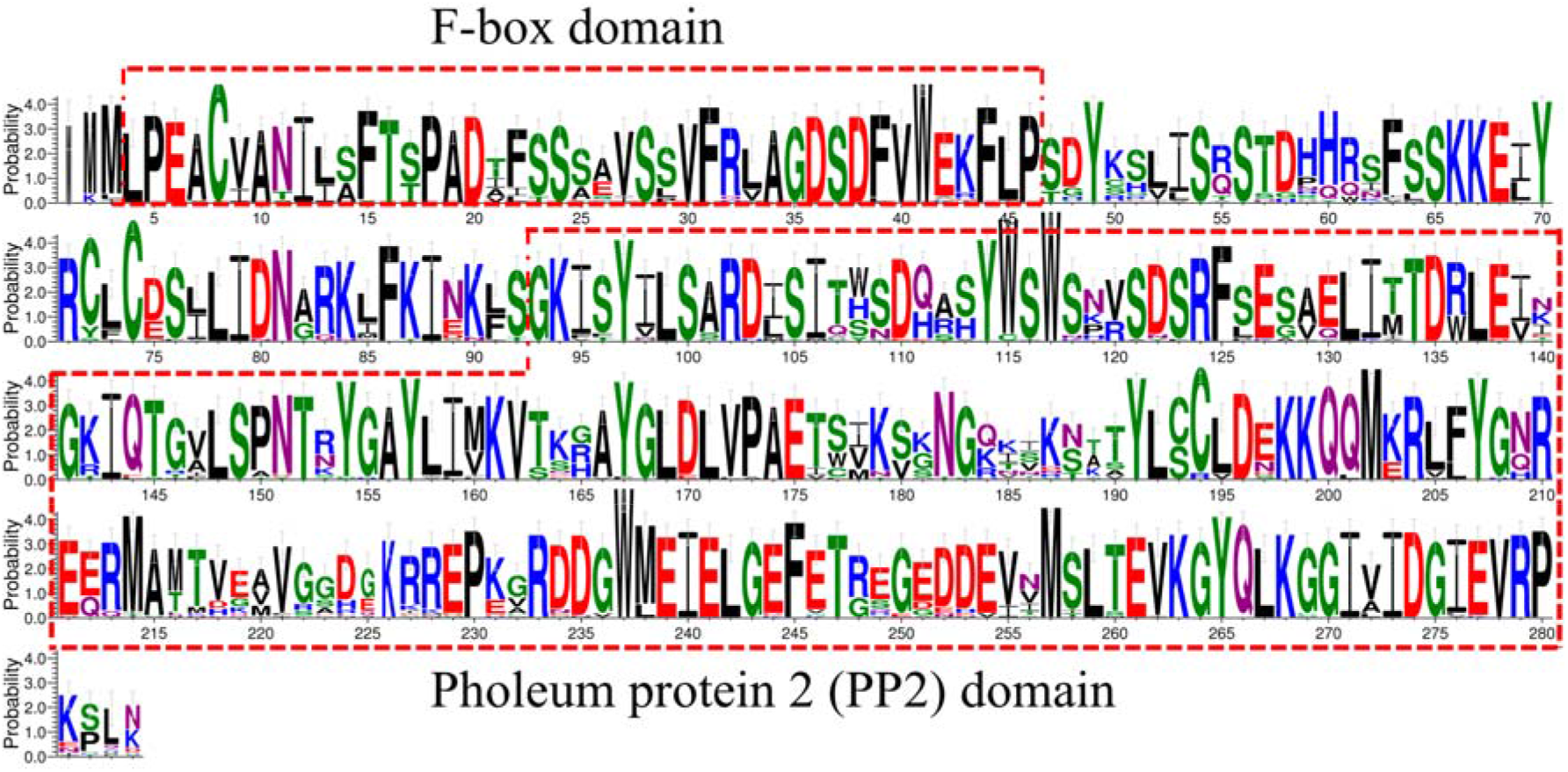
Different sequence conservation profiles in the PP2-B13 and its homologues in different plant species. Conservation plots were constructed using WEBLOGO. The y-axis represents the probability score. Y = 4 corresponds to 100% conservation. The predicted domains are highlighted in red boxes.

Lectins are proteins containing at least one non-catalytic domain which enables them to selectively recognize and bind to specific glycans that are either present in a free form or are part of glycoproteins and glycolipids and help the plants to sense the presence of pathogens; as a defense response they use a broad variety of lectin domains to interact with pathogens^58^. Additionally, Eggermont et al., (2017)^57^ showed that among lectin proteins, the amino acids responsible for carbohydrate binding are highly conserved. Furthermore, Jia et al (2015)^58^, showed that PP2-B11, (another member of the phloem lectin proteins^48^) is highly induced in response to salt treatment at both transcript and protein levels. They showed that PP2-B11 plays a positive role in response to salt stress.

In order to predict PP2-B13 interaction partners, we submitted the PP2-B13 amino acid sequence to the STRING database (version 11.0), which hypothetically determines protein– protein interactions based on computational prediction methods^59^. This returned several major players in innate immunity, specifically PBL1, RLP6 and RLP15, which are important defense proteins, as potential interaction partners (Fig. 9)^60–62^. RLPs are regarded major players in immune system in Arabidopsis^60–62^. STRING also predicted interactions of PP2-B13 with major zinc transporter proteins (ZIPs), which have role in biotic and abiotic stress responses^63^.

**Figure 9.**
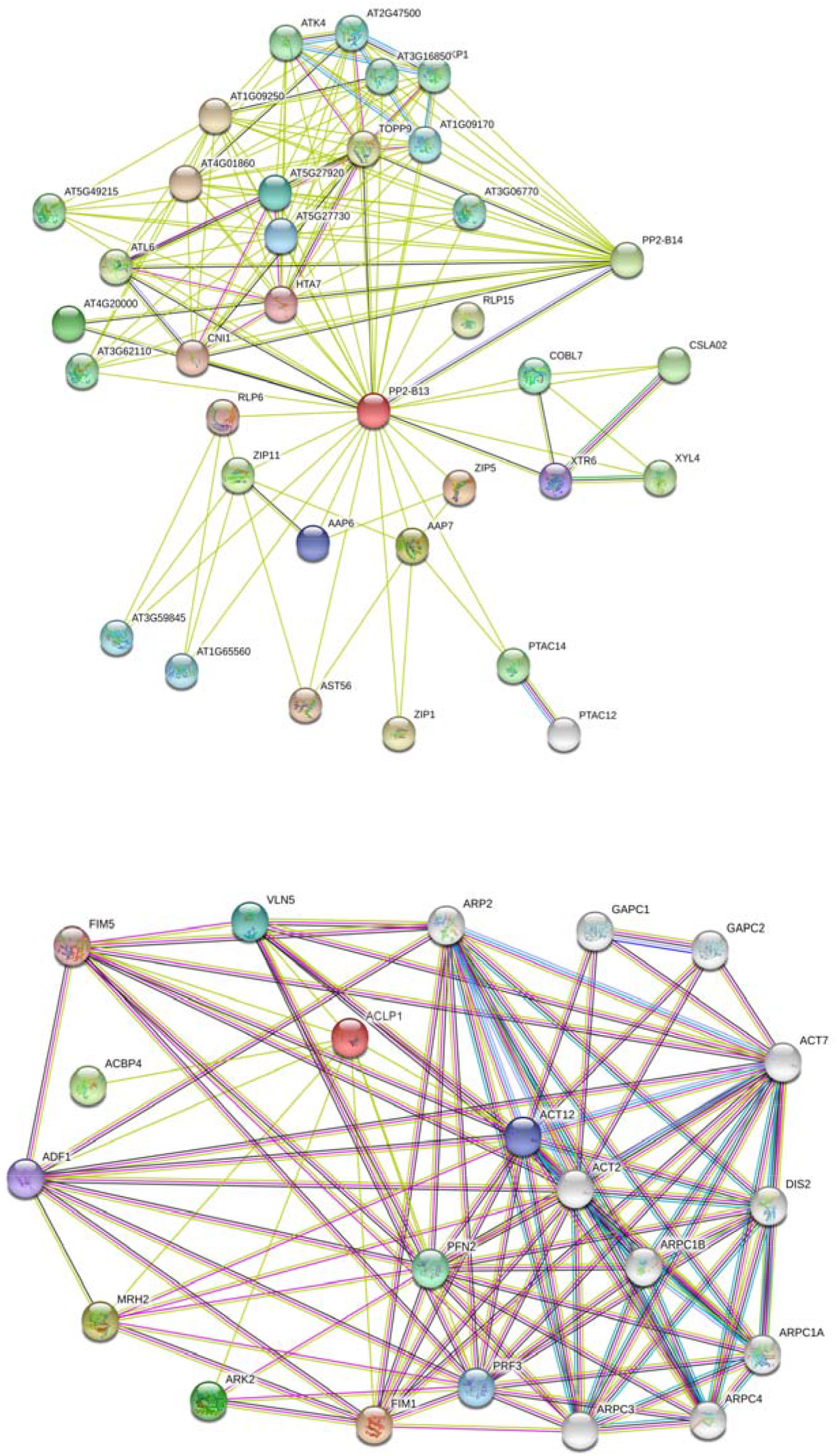
The protein-protein interaction (PPI) network of the PP2-B13 and ACLP1 proteins in *Arabidopsis thaliana* based on STRING 11.0. analysis with a confidence threshold score of 0.4 (Szklarczyk et al., 2015).^87^ Line colors indicate type of interaction used for the predicted associations: gene fusion (red), gene neighborhood (green), co-occurrence across genomes (blue), co-expression (black), experimental (purple), text mining (light green); association in curated databases (light blue). Line thickness represents the strength of data support. Proteins which have a known function in immune response are marked with dotted lines.

In the region of the chromosome 1 where PP2-B13 is located, there are many genes which are activated upon biotic or abiotic stresses, such as AT1G56280. The protein product of this gene is named drought-induced protein 19 (Di19), because its expression increases due to progressive drought stress^64^. Importantly, we have found that the *WRR4* gene (*AT1G56510*) is downstream of the *PP2-B13* (Supplementary Fig. S2). WRR4 is one of the most important defense gene in *Arabidopsis thaliana*^64–65^.

ACLP1 is an actin cross-linking protein of 397 amino acids. Raptor X^54^ predicted two Fascin motifs in the N -terminal and C-terminal domains (residues 18-70 and 229-318, respectively; Fig. 10; Supplementary Fig. S6). The conserved domain database at NCBI (https://www.ncbi.nlm.nih.gov/Structure/cdd/cdd.shtml) also identified two fascin domains in ACLP1. Fascins are a structurally unique and evolutionarily highly conserved group of actin cross-linking proteins. Fascins function in the organization of two major forms of actin-based structures: dynamic, cortical cell protrusions and cytoplasmic microfilament bundles^67–69^. For ACLP1 the sequence Logo was created. As shown in the Figure 10, there are several conserved regions in the ACLP1 and its homologues. Furthermore, a phylogenetic analysis supported high conservation of ACLP1-like proteins across different land plant species, suggesting similar function (Supplementary Fig. 9).

**Figure 10.**
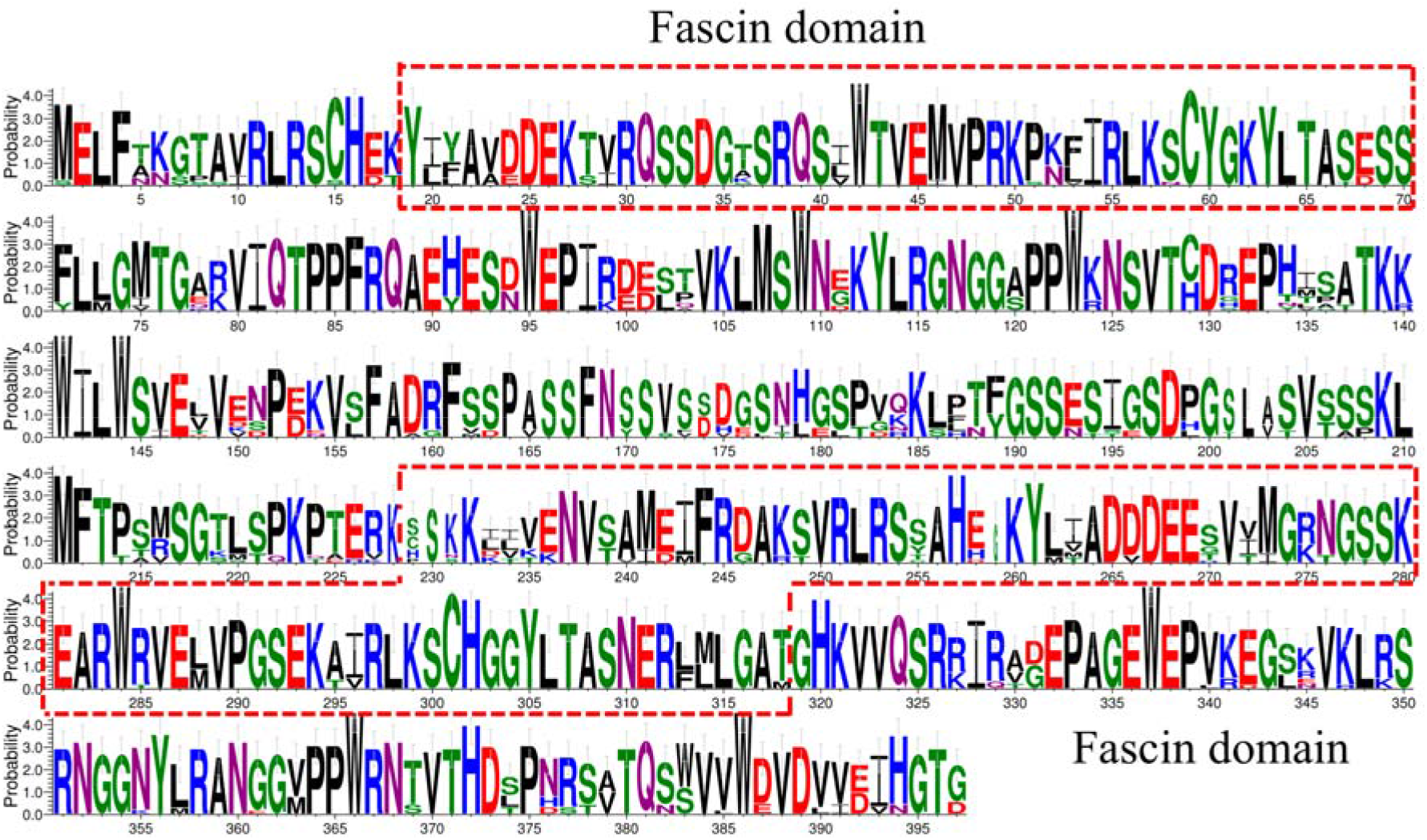
Different sequence conservation profiles in the ACLP1 and its homologues in different plant species. Conservation plots were constructed using WEBLOGO. The y-axis represents the probability score. Y = 4 corresponds to 100% conservation. The predicted domains are highlighted in red boxes.

MAMP perception changes actin arrangements and leads to cytoskeleton remodeling^70–71^. Cytoskeleton rapidly responds to biotic stresses to supports cellular fundamental processes^72–74^. Recently, Henty-Ridilla et al., (2014)^75^ confirmed that Actin depolymerizing factor 4 (ADF4) has an important role in defense response through cytoskeleton remodeling. They showed that the *adf4* mutant was unresponsive to a bacterial MAMP^76^. Using the STRING database (version 11.0), we predicted many actin related proteins including ADF4, ACT2, ACT12, PFN2, MRH2, ARK2 and ADF1 as putative interaction partner for ACLP1 (Fig. 9), further corroborating a potential role for ACLP in defense-related actin reorganization. It is noteworthy that downstream of the *ACLP1*, there is *PP2-A5* gene (Supplementary Fig. S3). The protein product of PP2-A5 gene is another member of the Phloem Protein 2 family. The role of PP2-A5 in defense response against insect is already confirmed^77^.

## Conclusions and outlook

We observed a strong (>100fold) and very rapid but transient induction of *PP2-B13* and *ACLP1* within 30 min of flg22 elicitor treatment (Fig. 2). Using a mutant approach, we provide evidence that loss-of-function mutations in *PP2-B13* and *ACLP1* can affect the early PTI responses including ET and ROS measurements (Fig. 5 and Fig. 6). We could show a defect in activation of ET production for *aclp1* plants and also attenuated ROS generation in *pp2-b13* plants in response to flg22 treatment. ROS accumulation is regarded as an early PTI event occurring a few minutes after *Pst* inoculation^1^. These findings suggest that these genes might have a function through interaction with PTI signaling pathways during bacterial infection. However, we cannot yet determine at what point of the MAMP signaling cascade the products of these genes function. Therefore, subsequent studies are needed to determine the relationship of these genes in MAMP recognition and other signaling cascades in innate immunity.

Furthermore, one important finding in this study is that our study reconfirm the importance of the chromosome 1 in innate immunity as there many resistant genes that their protein product have role in defense including ACLP1, Di19, PP2-A5, PP2-B13, WWR4 and VBF. Therefore, we suggest that in further studies, this region of the chromosome 1 should be evaluated in depth to identify more genes which have role in innate immunity.

In conclusion, based on what we have observed in different experiments, it can be concluded that PP2-B13 and ACLP1 have a role in innate immunity. It is likely that the protein products of these genes can have multiple functions in innate immunity in Arabidopsis. It has been previously reported that the genes which have a function in innate immunity in Arabidopsis can also have role in resistance against abiotic stress^78^. Hence, it will be interesting to see the response of these mutant plants upon abiotic stresses such as salinity, cold, and drought.

## Materials and Methods

### Plant material and growth conditions

All Arabidopsis genotypes were derived from the wild-type accession Columbia-0 (Col-0). The plants were grown as one plant per pot at 10 h photoperiod light at 21°C and 14 h dark at 18°C, with 60% humidity for 4 to 5 weeks, or were grown on plates containing Murashige and Skoog (MS) salts medium (Sigma, Aldrich), 1% sucrose, and 1% agar with a 16 h photoperiod. Seeds of the *sid2* mutant line were kindly provided by Jean-Pierre Métraux (University of Fribourg). The *fls2* mutant line was previously published^23^. *pp2-b13 (*AT1G56240; SALK_144757.54.50), and *aclp1* (AT1G69900; SALK_68692.47.55) were obtained from the Nottingham Arabidopsis Stock Centre (NASC).

### Peptide treatments

The peptides used as elicitors were flg22 (QRLSTGSRINSAKDDAAGLQIA), and *At*Pep1 (ATKVKAKQRGKEKVSSGRPGQHN). The peptides were ordered from EZBiolabs (EZBiolab Inc., IN, USA), dissolved in a BSA solution (containing 1 mg/mL bovine serum albumin and 0.1 M NaCl), and kept at −20°C. In order to prepare sterile seedlings, Arabidopsis seeds were washed with 99% ethanol supplemented with 0.5% Triton for 1 min, washed with 50% ethanol supplemented with 0.5% Triton for 1 min, then washed with 100% ethanol for 2 min. Seeds were sown on MS salt medium supplemented with 1% sucrose and 0.8% Phytagel (Sigma-Aldrich) at pH 5.7. Subsequently, the plates were stratified for 2 d at 4°C and germinated at 21°C under continuous light (MLR-350; Sanyo chamber). One day before treatment, the seedlings were moved from plates to ddH_2_O. One-week-old Arabidopsis seedlings were treated with *At*Pep1 and flg22 (1 μM) for 30 min. BSA solution was used for the mock-treated control.

### RNA isolation, Illumina sequencing and quality control

Total RNA was isolated from one-week-old Arabidopsis seedlings using the RNeasy Plant Mini Kit (Qiagen), according to the manufacturer’s protocol. Three individual biological replicates were used per condition. RNA purity, concentration, and integrity were determined via spectrophotometric measurement on a NanoDrop 2000 (Thermo-Scientific). Libraries were prepared using the RNA sample preparation kit (Illumina) according to the manufacturer’s instructions (Illumina). Libraries were sequenced on a HiSeq2000 instrument (Illumina) as 100 bp single-end reads. Sequencing quality of the fastq files from the RNA-Seq data was examined by FastQC software (version V0.10.1; http://www.bioinformatics.babraham.ac.uk/projects/fastqc/). Adapter sequences were clipped and low quality reads were either trimmed or removed.

### Mapping reads to the reference genome and analysis of differentially expressed genes (DEGs)

RNA-seq reads were aligned against the *A. thaliana* cDNA reference genome (TAIR10; (https://www.arabidopsis.org/). The reference genome index was constructed with Bowtie v2.2.3 and reads were aligned to the Arabidopsis reference genome using TopHat v2.0.12 with default parameters^46^. The resulting alignments were visualized using Integrative Genomics Viewer (IGV)^79^. To evaluate differentially expressed genes between elicitor-treated and control samples, we used the DESeq2 R package^80–81^. Genes with an adjusted *p*-value < 0.05 and a minimum two-fold change in expression were considered as differentially expressed.

### Selection of candidate genes

Because we were interested in genes not yet classified as related to immune response, we applied several filters: from the genes significantly up-regulated after 30 minutes of flg22 or AtPep1 treatment, we discarded those which had previously been reported as differentially regulated and implicated in biotic and abiotic stress response^23,40,41^. We selected a subset of 85 genes (Supplementary File 10) based on the following criteria: 1) high induction of transcription in response to both flg22 and AtPep1 treatments, 2) not present on Affymetrix ATH-22k microarray chips, 3) no published function or at least not connected to defense, and 4) not a member of a large gene family (in order to avoid potential functional redundancy). From this list, we eventually selected 20 genes as candidate genes for further analyses (Table 1) and ordered corresponding T-DNA insertion lines (http://signal.salk.edu/cgi-bin/tdnaexpress) from NASC (www.arabidopsis.info).

### Determination of gene expression by quantitative real-time RT-PCR analysis

Discs of leaves of four-week-old Arabidopsis plants were cut out using a sterile cork borer (d=7mm) and placed overnight in ddH_2_O in 5 cm Petri dish. Thereafter, the experiment started (time zero) with the addition of 1 μM flg22, dissolved in BSA solution (1 mg/mL bovine serum albumin and 0.1 M NaCl). BSA solution without flg22 was used for the mock-treated control., In order to produce a time course in response to flg22 treatment, the experiment was stopped after 30 min, 2 h and 6 h, Total RNA from leaves of four-week-old Arabidopsis plants was extracted using the NucleoSpin RNA plant extraction kit (Macherey-Nagel) and treated with rDNase according to the manufacturer’s extraction protocol. RNA quality of all samples was assessed using NanoDrop 2000 (Thermo-Scientific). To synthesize the cDNA, 10 ng of RNA was used with oligo (dT) primers and AMV reverse transcriptase and reverse transcription was performed according to the manufacturer’s instructions (Promega). Using a GeneAmp 7500 Sequence Detection System (Applied Biosystems), quantitative RT-PCR was performed in a 96-well format. The gene-specific primers used in this study are listed in Supplementary Table S2. Expression of *UBQ10* (AT4G05320), which has been validated for gene expression profiling upon flg22 treatment^82–84^, was used as the reference gene. Based on C_T_ values and normalization to *UBQ10* (*AT4G05320*) expression, the expression profile for each candidate gene was calculated using the qGene protocol^83–84^.

### Analysis of T-DNA insertion mutants

After grinding leaf material in liquid nitrogen, total DNA was extracted using EDM-Buffer (200 mM Tris pH7.5; 250 mM NaCl, 25 mM EDTA; 0.5% SDS). Putative T-DNA insertion mutants were genotyped by PCR. We designed gene specific primer pairs LP and RP based on the predicted genomic sequence surrounding the T-DNA insertion (Supplementary Table S2). The plants were considered homozygous mutants if there was a PCR product with T-DNA-specific border primers LP/ LBa1 but not with the LP/RP primers. (Table 1). We obtained T-DNA insertion mutants of six single homozygous lines bearing a disruption in the gene, including AT1G56240 (*PP2-B13*) and AT1G69900 (*ACLP1*) (Table 1).

### RT-PCR experiment

For total RNA extraction, samples of leaf tissue from 4-week-old Arabidopsis including wild type plants (Col0), *pp2-b13*, and *aclp1*were harvested into liquid nitrogen and were grounded with the sterile mortar and pestle. The NucleoSpin RNA Prep Kit (BioFACT™, South Korea) was used for RNA extraction according to the manufacturer’s instructions and DNase-treated.

Reverse transcription was performed at 50°C for 45 minutes using total RNA, a reverse transcriptase (BioFACT™, South Korea) and an oligo (dT)20 primer (BioFact, South Korea) supplemented with 0.5ul RNase inhibitor (BioFACT™, South Korea) and according to the manufacturer’s instructions. To ensure specificity and accuracy of each primer and to design the highly specific primers for *PP2-B13* and *ACLP1* transcripts, the oligonucleotide primers were designed by AtRTPrimer program^85^ which exclusively determine specific primers for each individual transcript in Arabidopsis. The housekeeping gene *ACTIN2* was used as a positive control for each PCR. The primers for *ACTIN 2* transcript were used as described previously^86^. Primers that were used in these experiments are listed in Supplementary Table S2.

### Bacterial growth assay

*Pseudomonas syringae* pathovar *tomato* mutant *hrcC-* (deficient in type three effector secretion system)^87,88^; was grown in 20 ml liquid YEB medium supplemented with 50 μg/ml Rifampicin on a shaker at 28°C overnight. Infection assay and counting the bacterial titer was done as described previously^89^ with a bacterial suspension at OD_600_ = 0.0002. Leaves of 4-5-week-old Arabidopsis plants were infiltrated using a syringe. The *sid2-2* mutant plants, which are incapable of accumulating salicylic acid^90^, were used as a positive control. Mock-infected plants were similarly treated with infiltration buffer.

### Measurement of ethylene production

Leaf material of Arabidopsis plants was cut into discs of 10 mm^2^ using a sterile cork borer, at the end of the light period. After mixing leaf strips from several plants, six leaf strips were placed together in a 6 ml glass vial containing 0.5 ml of ddH_2_O. Vials with leaf strips were incubated overnight in the dark in a short-day room (16 h dark / 8 h light). The following day (approximately after 16 h), elicitor peptide was added to the desired final concentration (1 μM) and vials were closed with air-tight rubber septa and put in the short-day room. ET accumulating in the free air space was measured by gas chromatography (GC-14A Shimadzu) after 4 h of incubation with or without elicitor.

### ROS measurement

Using a sterile cork borer, leaf discs of approximately 10 mm^2^ were cut from several plants. One leaf disc per well was left floating overnight in darkness in 96-well plates (LIA White, Greiner Bio-One) on 100 μl ddH_2_O at 18°C. Horseradish peroxidase (1 μg/ml final concentration), luminol (100 μM final concentration) and elicitor peptide (1 μM final concentration) were added to the wells. Using a plate reader (MicroLumat LB96P, Berthold Technologies) light emission of oxidized luminol in the presence of peroxidase was determined over 30 min, starting from addition of the elicitor.

### Immunoblot analysis

150 mg of leaf material from 4-5-week-old Arabidopsis plants was shock-frozen and ground in liquid nitrogen. 200 μl Läemmli buffer containing 50 mM β-mercaptoethanol was added and the ground homogenate was further mixed by vortexing. Proteins were denatured by boiling for 10 min at 95 °C. Debris was pelleted by centrifugation for 5 min at 13,000 rpm. Total proteins were separated by electrophoresis in 7% SDS-polyacrylamide gels and electrophoretically transferred to a polyvinylidene fluoride membrane according to the manufacturer’s protocol (Bio-Rad). Transferred proteins were detected with Ponceau-S. The abundance of FLS2 receptor was analyzed by immunoblot and immunodetection with anti-FLS2-specific antibodies as previously described^91^.

### Phylogenetic analysis and comparison consensus of the amino acid sequences

Protein sequences BB2-B13 and ACLP1 were used as queries using BLASTP^92^ (Basic Local Alignment Search Tool) search to identify the most similar proteins in *A. thaliana* and diverse land plants. We applied a cutoff of 70%≤ sequence identity on the top hit of the BLASTP search for BB2-B13 and ACLP1 and their orthologous and prologues were identified. Protein sequences with more than 70% sequence identity were downloaded from the NCBI database and multiple alignment analysis performed based on the ClustalW software^93^. Phylogenetic analyses and graphical representation were carried out using MEGA software (Molecular Evolutionary Genetics Analysis) version 6.0^94^. A neighbor-joining phylogenetic tree was constructed after 1,000 iterations of bootstrapping of the aligned sequences. All branches with bootstrap values <60% were collapsed. To compare consensus of the amino acid sequences, sequence logos were generated using WebLogo (http://www.weblogo.berkeleky.edu/), using the ClustalW alignment as input.

## Abbreviations

DAMPs: Damage-associated molecular patterns
DEGs: differentially expressed genes
MAMPs: microbe-associated molecular patterns
NGS: next generation sequencing
PRR: pattern-recognition receptor
PEPR: AtPep-receptor
PP2: phloem protein 2
PTI: pattern-triggered immunity

## Acknowledgments

The authors would like to sincerely acknowledge Dr Sebastian Merker (University of Basel) for his skillful technical advices and very helpful discussion. We would like to sincerely thanks Dr Jonathan Seguin (University of Strasburg, France) for his helpful comments and advice for data analysis and discussion about the results. The authors wish to thank Dr. Delphine Chinchilla (University of Basel, Switzerland) for her helpful comments and useful discussions. We are grateful to Dominik Klauser (University of Basel; Syngenta Foundation for Sustainable Agriculture, Switzerland) for helping us to set up the experiments. We are sincerely thankful to Peter Palukaitis (Seoul Women’s University, South Korea) for his critical review and proofreading of the manuscript. We are thankful to the GDC center at the University of ETH Zurich for their helpful comments. Special thanks to the researchers at the Faculty of Sciences and Biotechnology at Shahid Beheshti University (Tehran, Iran) for very helpful discussion and useful comments.

## Author Contributions

T.B., M.S., and C.B. conceived and designed the work. T.B. conceptualized the research work. M.S. and C.B. performed sequence analysis and data analysis. M.S. performed most of the data mining and experiments. M.S. wrote the original draft of the manuscript. T.B. and C.B. revised the manuscript. All authors read and approved the final manuscript.

## Disclosure of Potential Conflicts of Interest

No potential conflicts of interest were disclosed.

## Funding

This research was supported by Freiwillige Akademische Gesellschaft (FAG) award, Basel, Switzerland [15_3_2015], Niklaus und Bertha Burckhardt-Buergin-Stiftung Foundation, Basel, Switzerland [22_3_2016] and the Shahid Beheshti University research fund, Tehran, Iran.

## Supplementary Information

**Supplementary File S1.** List of all genes in response to flg22 treatment compared to the BSA treatment

**Supplementary File S2.** List of all genes in response to AtPep1 treatment compared to the BSA treatment

**Supplementary File S3.** List of top up-regulated DEGs in response to flg22 treatment compared to the BSA treatment

**Supplementary File S4.** List of top up-regulated DEGs in response to AtPep1 treatment compared to the BSA treatment

**Supplementary File S5.** List of top down-regulated DEGs in response to flg22 treatment compared to the BSA treatment

**Supplementary File S6.** List of top down-regulated DEGs in response to AtPep1 treatment compared to the BSA treatment

**Supplementary File S7.** List of the genes which their expression exclusively identified in the RNA-seq transcriptional profiling analysis in response to flg22 treatment and are not present in Affymetrix ATH1 GeneChip

**Supplementary File S8.** List of the up-regulated DEGs with fold change cutoff in response to flg22 treatment compared to the BSA treatment that are present in both RNA-seq experiment analysis and ATH1 affymetirx genechip

**Supplementary File S9.** List of the up-regulated DEGs with fold change cutoff in response to flg22 treatment compared to the BSA treatment that are exclusively present in RNA-seq experiment analysis

**Supplementary File S10.** List of 85 selected genes for subsequent study

**Supplementary File S11.** List of DEGs exclusively up-regulated in response to flg22 treatment compared to the BSA treatment

**Supplementary File S12.** List of DEGs exclusively up-regulated in response to AtPep1 treatment compared to the BSA treatment

**Supplementary File S13.** List of DEGs exclusively down-regulated in response to flg22 treatment compared to the BSA treatment

**Supplementary File S14.** List of DEGs exclusively down-regulated in response to AtPep1 treatment compared to the BSA treatment

**Supplementary File S15.** List of up-regulated DEGs in response to flg22 treatment compared to the AtPep1 treatment

**Supplementary File S16.** List of down-regulated DEGs in response to flg22 treatment compared to the AtPep1 treatment

## Supplementary Information

**Supplementary Table S1.** Summary of Illumina sequencing data and mapped reads of *Arabidopsis thaliana* wild-type (Col-0) under BSA, flg22 and AtPep1 treatments.

|  | treatment |  |  |  |  |  |  |  |  |
| --- | --- | --- | --- | --- | --- | --- | --- | --- | --- |
|  | BSA |  |  | flg22 |  |  | AtPep1 |  |  |
|  | Repeat 1 | Repeat 2 | Repeat 3 | Repeat 1 | Repeat 2 | Repeat 3 | Repeat 1 | Repeat 2 | Repeat 3 |
| **Total clean read** | 1239300 | 12961695 | 14519871 | 17431550 | 9827458 | 14682108 | 12463709 | 10905959 | 12881404 |
| **Mapped reads of input** | 11934302 | 12646663 | 14157965 | 16833273 | 9581401 | 14350429 | 12162285 | 10620059 | 12538057 |
| **Read mapping rate of input (percent)** | 96.3% | 97.6% | 97.5% | 96.6% | 97.5% | 97.7% | 97.6% | 97.4% | 97.3% |
| **Unmapped reads** | 210314 | 227871 | 273224 | 310845 | 170924 | 25657 | 22303 | 189382 | 222259 |
| **Unmapped reads (percent)** | 1.8% | 1.8% | 1.9% | 1.8% | 1.8% | 1.8% | 1.8% | 1.8% | 1.8% |

**Supplementary Figure S1.**
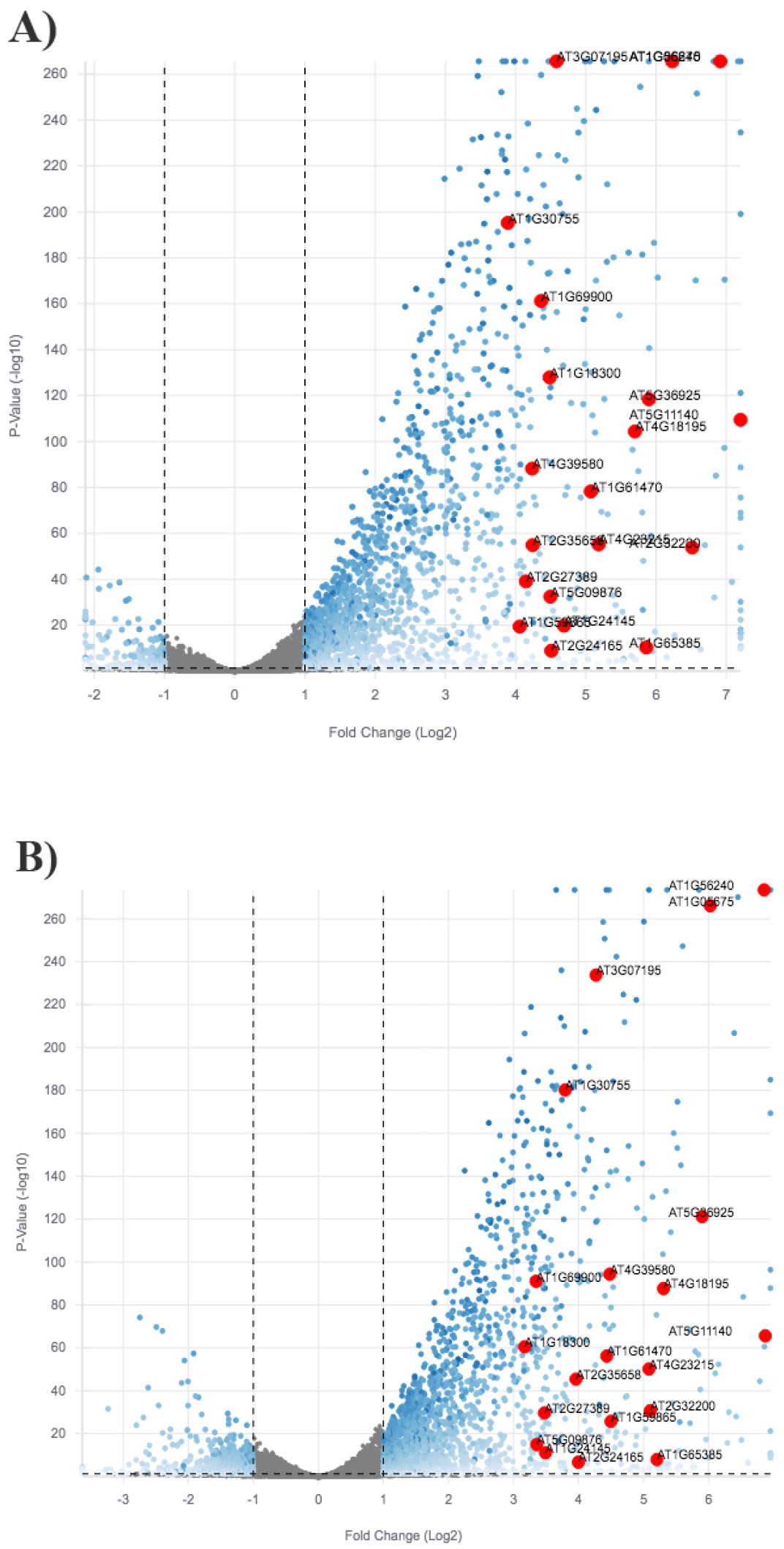
Volcano plot of gene expression in the seedling of Arabidopsis in response to flg22 treatment (**A**) and AtPep1 treatment (**B**). Blue dots correspond to significantly up- and down-regulated DEGs, while non-DEGs are in grey color. R

**Figure S2.**
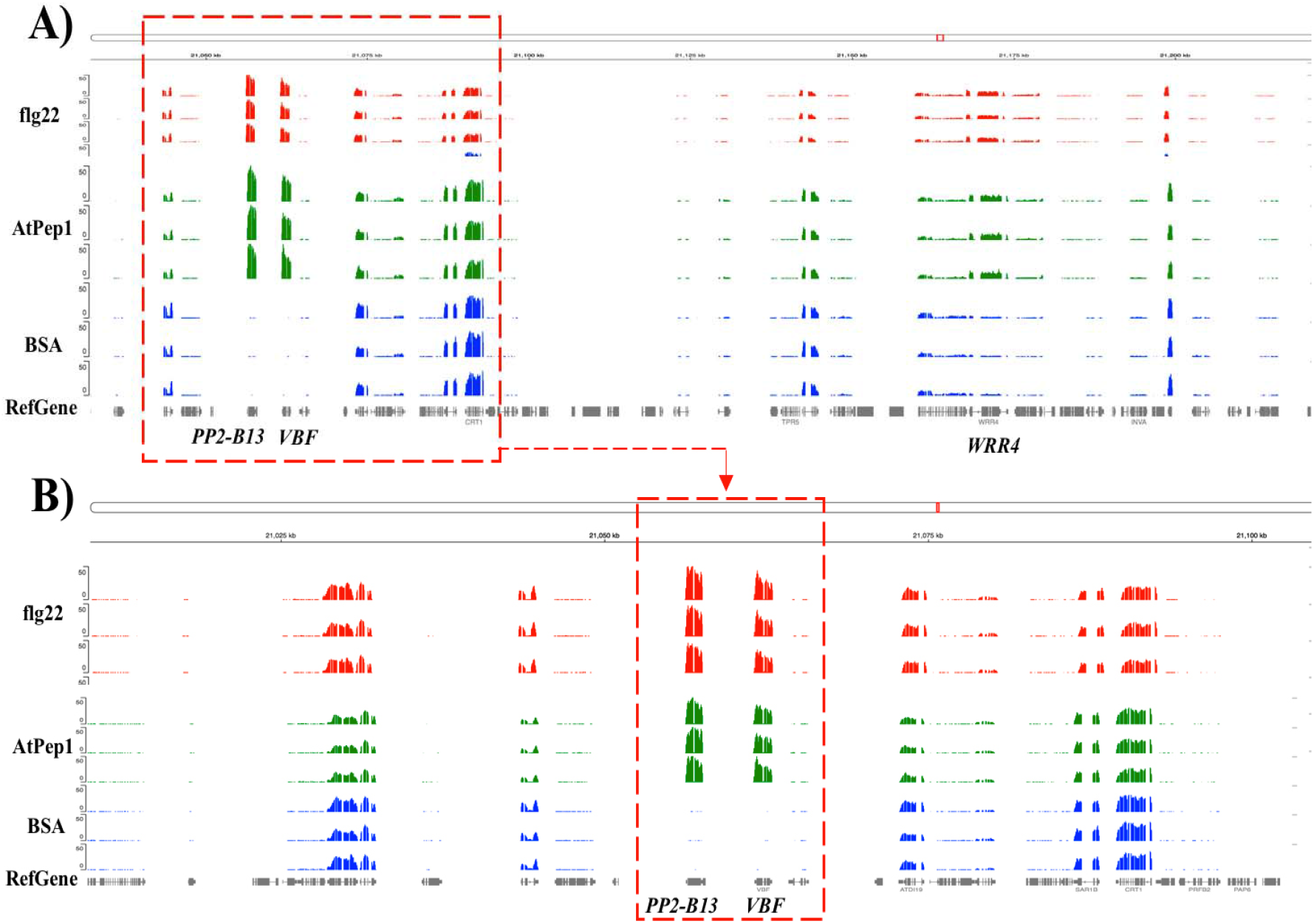
Integrative genomics viewer (IGV) visualization of alignments and coverage of the Illumina reads at the PP2-B13 locus. Coverage depth graphs represent transcript abundance. (**A**) Overlaid depth graphs. (**B**) Zoomed in view of A. In the graph *PP2-B13*, *VBF* and *WRR4* genes are illustrated.

**Figure S3.**
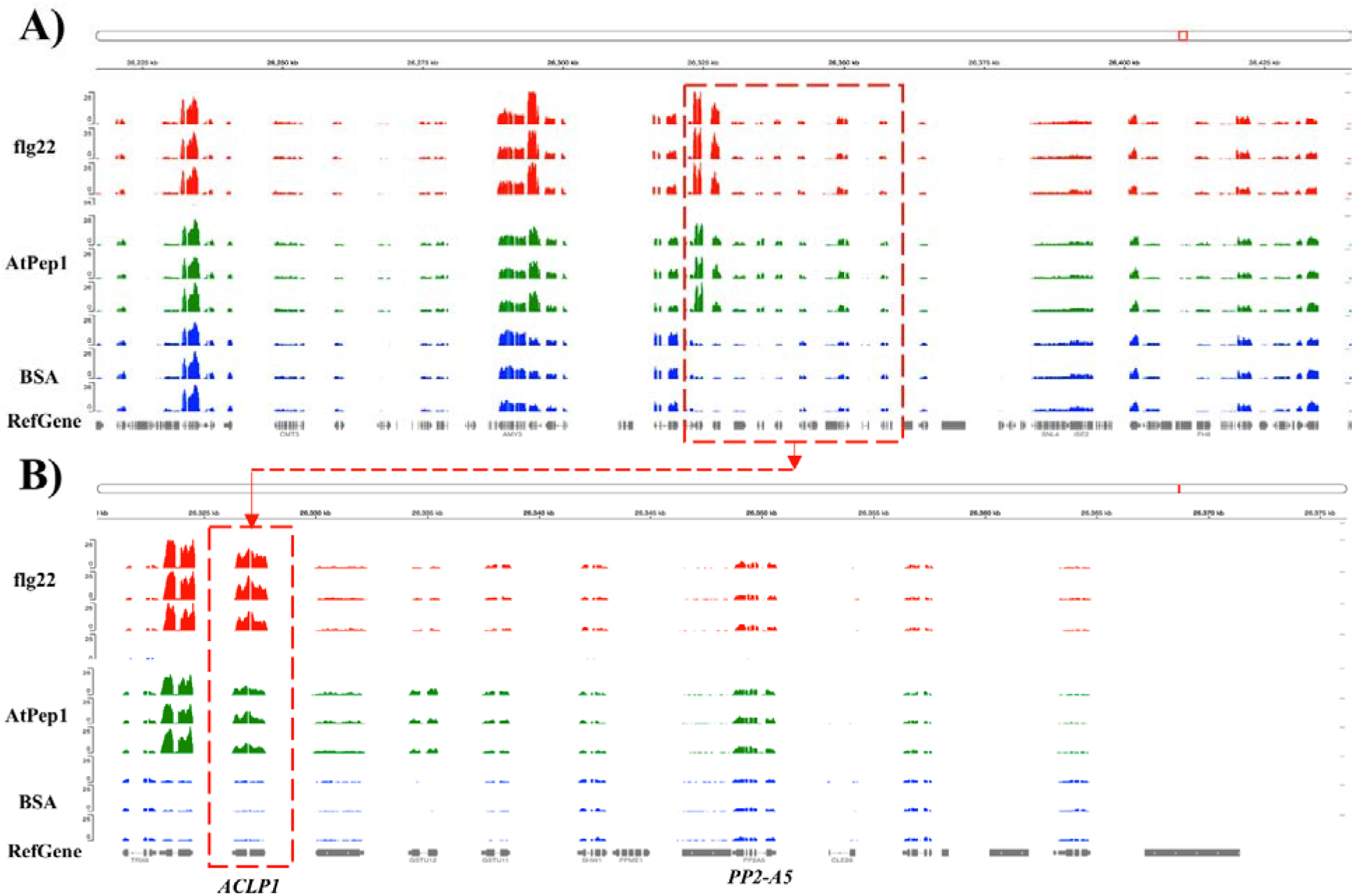
Integrative genomics viewer (IGV) visualization of alignments and coverage of the Illumina reads at the ACLP1 locus. (**A**) Overlaid depth graphs. (**B**) Zoomed in view of A. In the graph *ACLP1* and *PP2-A5* genes are illustrated.

**Supplementary Figure S4.**
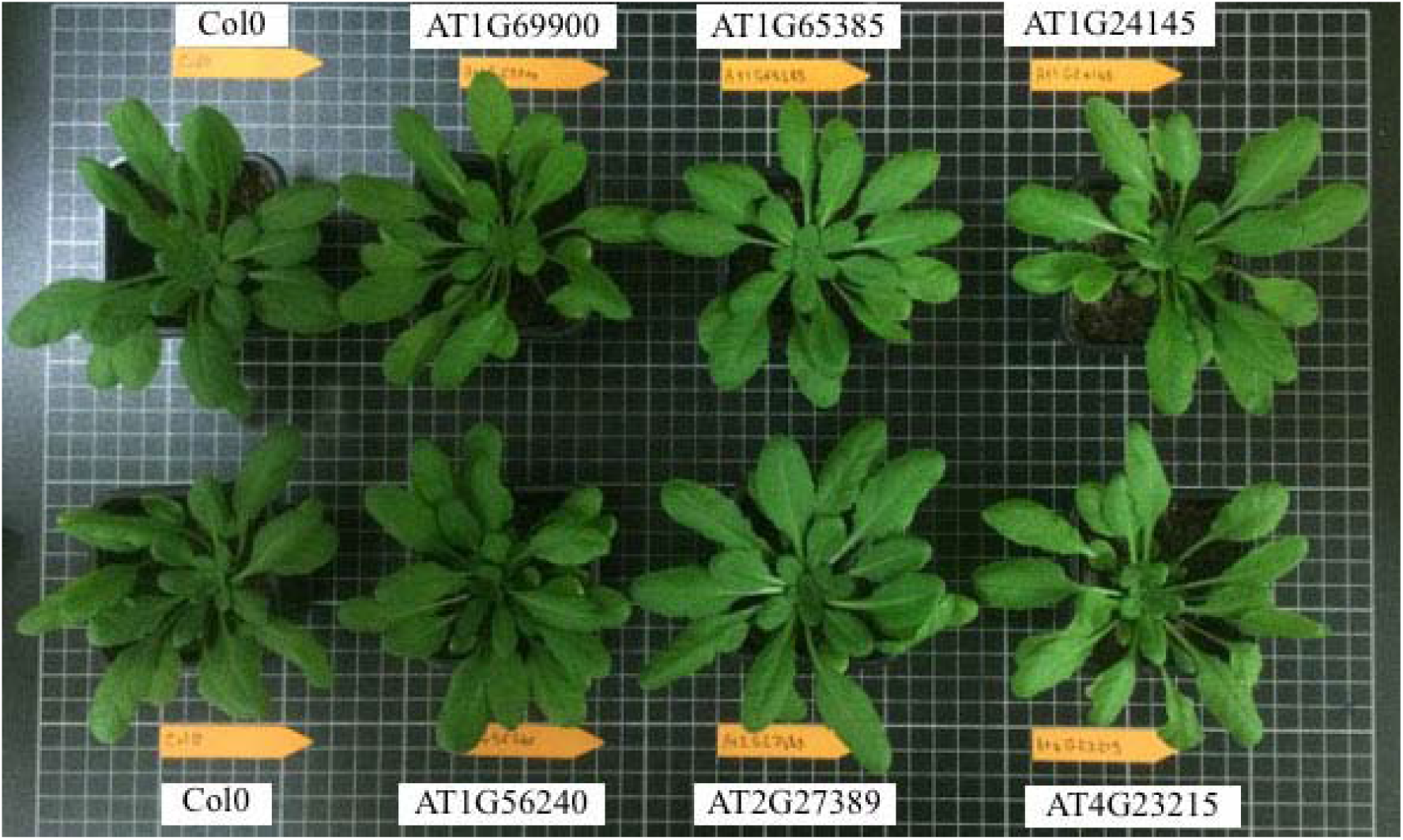
Phenotype of five-week old Arabidopsis plants. Plant were grown under short-day conditions (ten hours light at 21°C and 14 hours dark at 18°C, with 60% humidity).

**Supplementary Figure S5.**
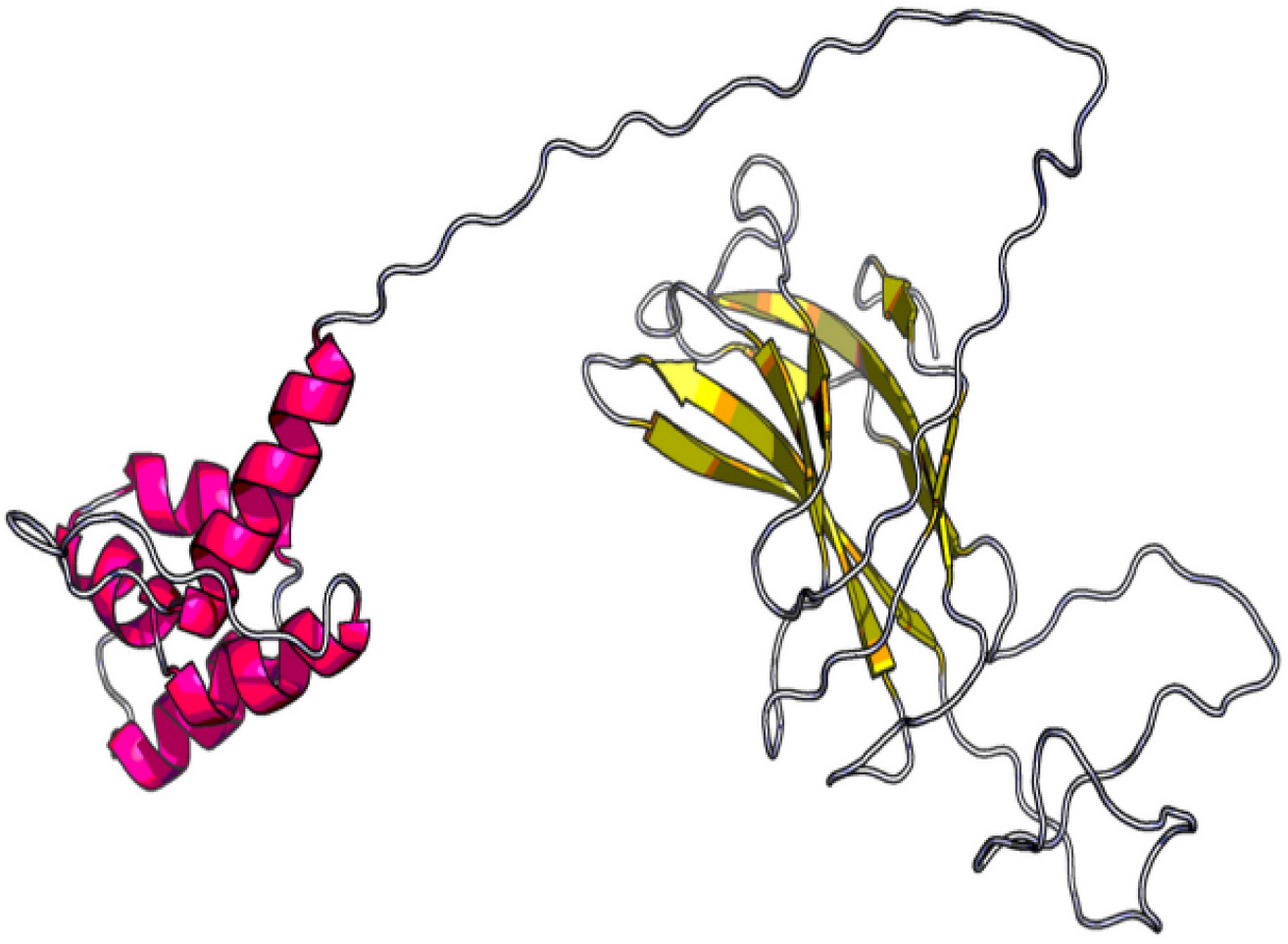
Structure of PP2-B13 protein determined by Raptor X (Källberg *et al*., 2012).

**Supplementary Figure S6.**
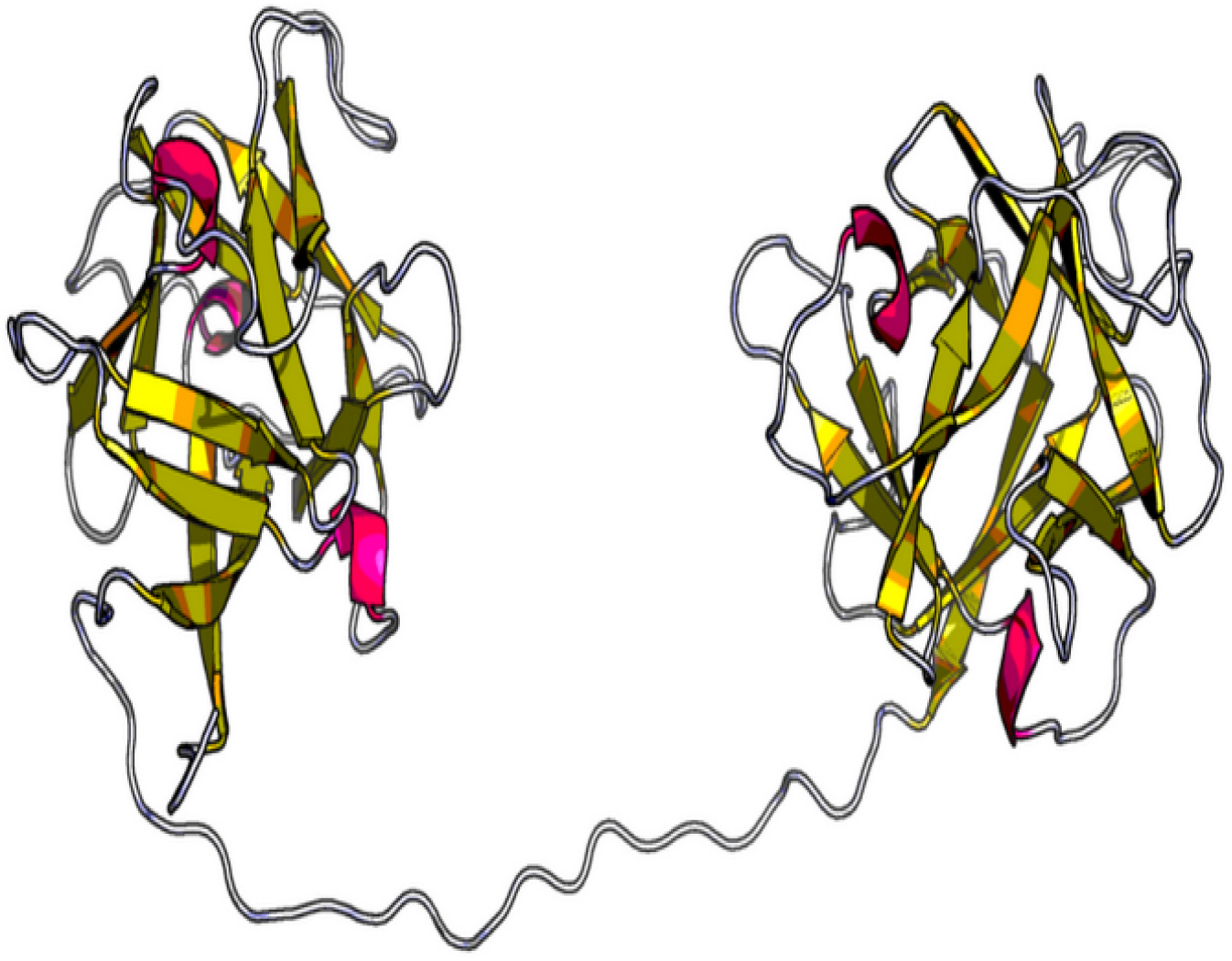
Structure of ACLP1 protein determined by Raptor X (Källberg *et al*., 2012).

**Supplementary Figure S7.**
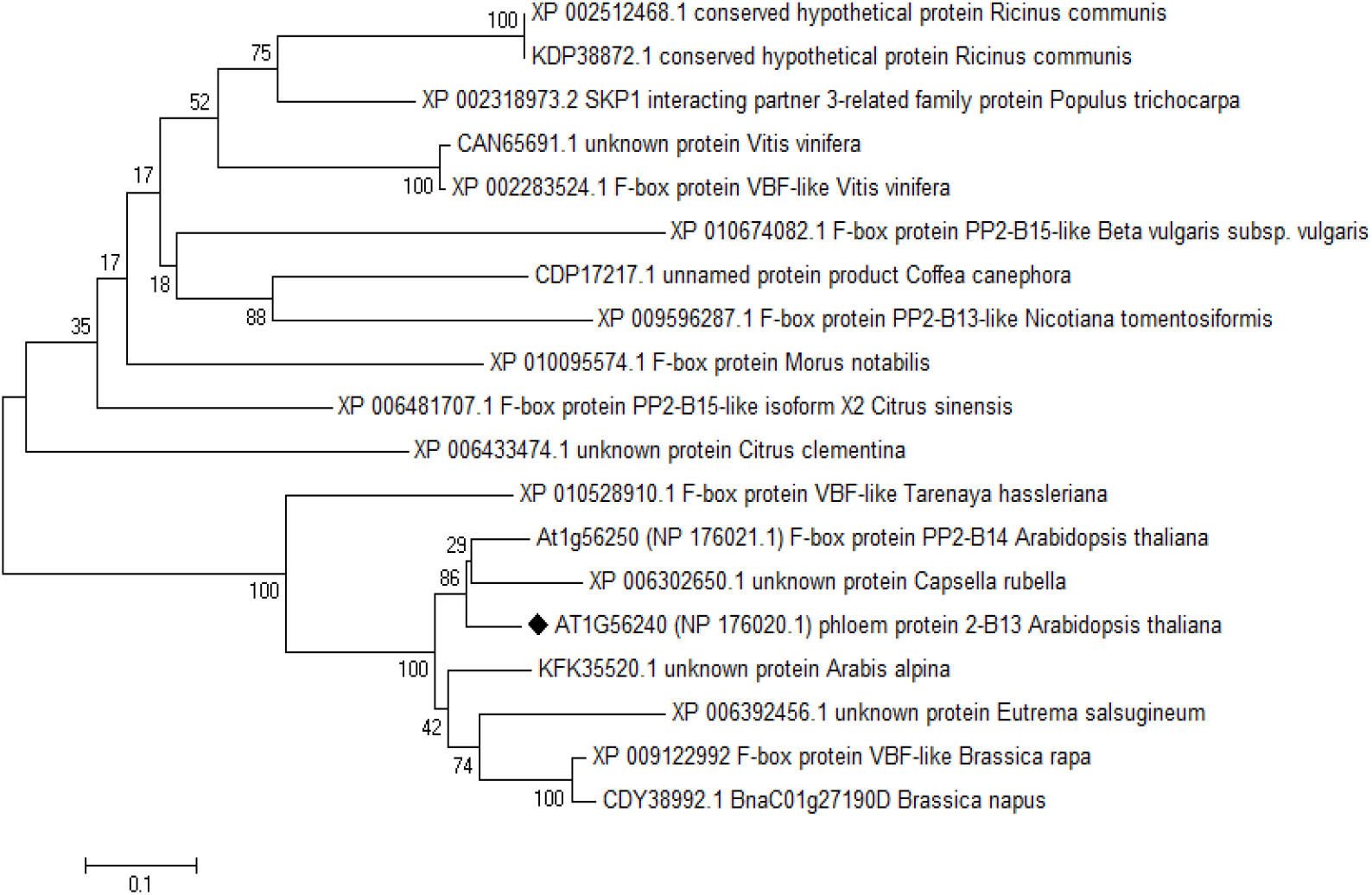
Phylogenetic analysis from the sequences of PP2-B13 protein in *Arabidopsis thaliana* and 15 representative land plants. The species indicated are *Ricinus communis*, *Populus trichocarpa*, *Vitis vinifera*, *Beta vulgaris*, *Coffea canephora*, *Nicotiana tomentosiformis*, *Morus notabilis*, *Citrus sinensis*, *Citrus clementine*, *Tarenaya hassleriana*, *Arabidopsis thaliana, Capsella rubella*, *Arabis alpina*, *Eutrema salsugineum*, *Brassica rapa* and *Brassica napus*. PP2-B13 protein in *Arabidopsis thaliana* was labelled. Sequences for comparisons were obtained from GenBank. The accession numbers and protein names (if available) are given. Analysis was done by maximum likelihood method implemented in MEGA6 (Molecular Evolutionary Genetics Analysis) version 6.0.

**Supplementary Figure S8.**
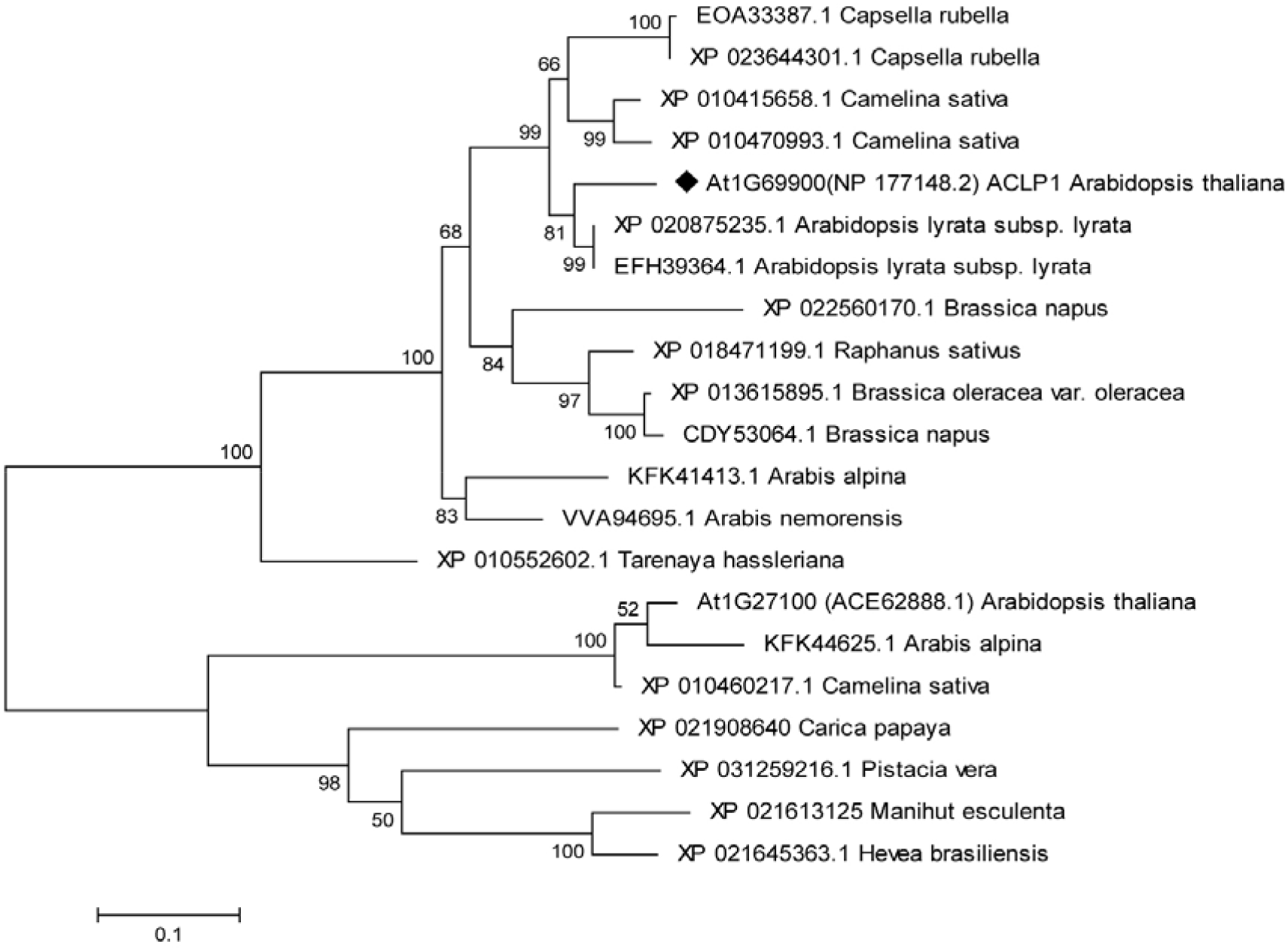
Phylogenetic analysis from the sequences of ACLP1 protein in *Arabidopsis thaliana* and 14 representative land plants. The species indicated are *Capsella rubella*, *Camelina sativa*, *Arabidopsis thaliana, Arabidopsis lyrata*, *Brassica napus*, *Raphanus sativus*, *Brassica oleracea*, *Arabis alpina*, *Arabis nemorensis*, *Tarenaya hassleriana*, *Carica papaya*, *Pistacia vera*, *Manihut esculenta* and *Hevea brasiliensis*. ACLP1 protein in *Arabidopsis thaliana* was labelled. Sequences for comparisons were obtained from GenBank. The accession numbers and protein names (if available) are given. Analysis was done by maximum likelihood method implemented in MEGA6 (Molecular Evolutionary Genetics Analysis) version 6.0.

**Supplementary Table S2.**
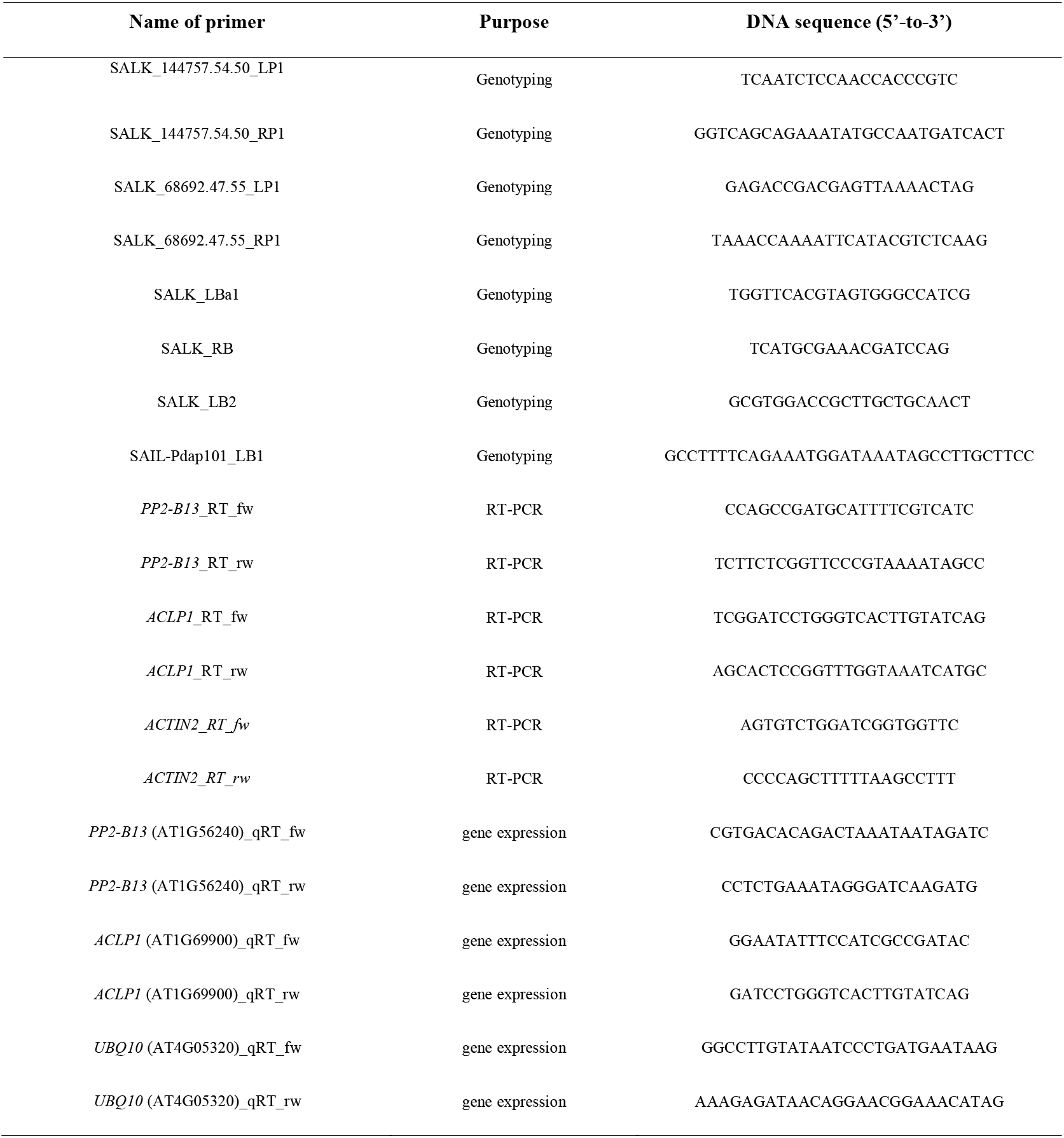
List of the oligonucleotide primers which were used in this study.

## Notes

https://www.ebi.ac.uk/arrayexpress/experiments/E-MTAB-9838/

### Competing Interest Statement

The authors have declared no competing interest.

